# Plasma membrane phosphatidylinositol-4-phosphate is not necessary for *Candida albicans* viability, yet is key for cell wall integrity and systemic infection

**DOI:** 10.1101/2021.12.31.474627

**Authors:** Rocio Garcia-Rodas, Hayet Labbaoui, François Orange, Norma Solis, Oscar Zaragoza, Scott Filler, Martine Bassilana, Robert A. Arkowitz

## Abstract

Phosphatidylinositol phosphates are key phospholipids with a range of regulatory roles, including membrane trafficking and cell polarity. Phosphatidylinositol-4-phosphate [PI(4)P] at the Golgi is required for the budding to filamentous growth transition in the human pathogenic fungus *Candida albicans*, however the role of plasma membrane PI(4)P is unclear. We have investigated the importance of this phospholipid in *C. albicans* growth, stress response, and virulence by generating mutant strains with decreased levels of plasma membrane PI(4)P, *via* deletion of components of the PI-4-kinase complex, *i.e.* Efr3, Ypp1 and Stt4. The amount of plasma membrane PI(4)P in the *efr3Δ/Δ* and *ypp1Δ/Δ* mutant was ∼60% and ∼40% of the wild-type strain, respectively, whereas it was nearly undetectable in the *stt4Δ/Δ* mutant. All three mutants had reduced plasma membrane phosphatidylserine (PS). Although these mutants had normal yeast phase growth, they were defective in filamentous growth, exhibited defects in cell wall integrity and had an increased exposure of cell wall β(1,3)-glucan, yet they induced a range of hyphal specific genes. In a mouse model of hematogenously disseminated candidiasis, fungal plasma membrane PI(4)P levels directly correlated with virulence; the *efr3Δ/Δ* had wild-type virulence, the *ypp1Δ/Δ* mutant had attenuated virulence and the *stt4Δ/Δ* mutant caused no lethality. In the mouse model of orpharyngeal candidiasis, only the *ypp1Δ/Δ* mutant had reduced virulence, indicating that plasma membrane PI(4)P is less important for proliferation in the oropharynx. Collectively, these results demonstrate that plasma membrane PI(4)P levels play a central role in filamentation, cell wall integrity and virulence in *C. albicans*.

**Importance:** While the PI-4-kinases Pik1 and Stt4 both produce PI(4)P, the former generates PI(4)P at the Golgi and the latter at the plasma membrane and these two pools are functionally distinct. To address the importance of plasma membrane PI(4)P in *Candida albicans,* we have generated deletion mutants of the three putative plasma membrane PI-4-kinase complex components and quantified the levels of plasma membrane PI(4)P in each of these strains. Our work reveals that this phosphatidylinositol phosphate is specifically critical for the yeast-to-hyphal transition, cell wall integrity and virulence in a mouse systemic infection model. The significance of this work is in identifying a plasma membrane phospholipid that has an infection specific role, which is attributed to the loss of plasma membrane PI(4)P resulting in β(1,3)-glucan unmasking.

## Introduction

Phosphatidylinositol phosphates are minor components of cellular membranes that play an essential role during polarized growth. In particular, phosphatidylinositol-4-phosphate [PI(4)P] is found predominantly at the Golgi and plasma membranes, generated from the precursor PI which is synthesized at the cytosolic face of the ER (1). Type III PI4-kinases are found in virtually all eukaryotes, with Pik1 in fungi homologous to mammalian type III*β* PI4-kinases and Stt4 to mammalian type IIIα PI4-kinases (2, 3). While all fungi appear to have Stt4 orthologs, Pik1 orthologs are absent in some of them, including *Aspergillus nidulans* and *Cryptococcus neoformans* (4, 5) and it has been suggested that Stt4 can carry out Pik1 essential function in these fungi. Plasma membrane PI(4)P generated by Stt4, has been implicated in the control of membrane trafficking, lipid exchange, cell signaling, cytoskeleton organization and cytokinesis. Stt4 type IIIα PI4-kinases are essential for viability in most fungi including *Saccharomyces cerevisiae* and *Schizosaccharomyces pombe* (6–9), however *stt4* mutants can be rescued by osmoremediation (6, 9).

*Candida albicans* is a major human fungal opportunistic pathogen that grows in both yeast and filamentous forms. The morphological transition between yeast and filamentous growth is important for its virulence and can be triggered by a range of stimuli (10–12). Using mutants in which the levels of phosphatidylinositol phosphates can be reduced revealed that Golgi PI(4)P and plasma membrane PI(4,5)P_2_ are critical for this transition (13, 14). Mutants with reduced levels of the plasma membrane PI-4-kinase Stt4, can nonetheless form germ tubes and grow invasively (14), raising the question as to whether this pool of PI(4)P is critical for the yeast-to-hyphal transition. In *C. albicans* (13), as in *S. cerevisiae* (15, 16), the Golgi and plasma membrane pools of PI(4)P are functionally distinct, *i.e.* Golgi PI(4)P does not substantially contribute to plasma membrane PI(4)P. A previous study suggested that the plasma membrane type IIIα PI4-kinase Stt4 was not essential for viability in *C. albicans* (17), raising the possibility that plasma membrane PI(4)P may be dispensable.

In yeast and mammalian cells, Stt4 is part of a complex comprised of the membrane protein Efr3 and a scaffold protein Ypp1 (TTC7 in mammals). Efr3 and Ypp1 are required for both targeting of Stt4 to the plasma membrane and PI-4-kinase activity (18–20). Specifically, Efr3 is critical for the plasma membrane association of the PI-4-kinase complex and Ypp1/TTC7 has been shown to bind directly both Efr3 and Stt4 (18, 19), suggesting a scaffolding function. In *S. cerevisiae*, both Ypp1 and Efr3 are essential for viability (18, 21, 22), whereas in *S. pombe* only Ypp1 is essential for viability (8). In conditional *ypp1* and *efr3 S. cerevisiae* mutants, cellular PI(4)P levels were reduced approximately two-fold (18, 20, 22).

To investigate the importance of plasma membrane PI(4)P in *C. albicans* hyphal growth and virulence we have generated deletion mutants of *efr3* and *ypp1*, in addition to *stt4*. These mutants were all viable and had different levels of plasma membrane PI(4)P. Surprisingly, *C. albicans* cells with little to no plasma membrane PI(4)P remain viable and can proliferate by budding growth. Furthermore, our results indicate that plasma membrane PI(4)P is critical for hematogenously disseminated candidiasis (HDC) with a *ypp1* mutant exhibiting reduced virulence and an *stt4* mutant causing no lethality, consistent with their plasma membrane PI(4)P levels. Our analyses of the cell wall suggest that the dramatic reduction in virulence in HDC is, in part, due to an unmasking of cell surface β(1,3)-glucan.

## Results

### The non-essential Stt4 PI-4-kinase complex is critical for invasive filamentous growth and cell wall integrity

Previously, we have generated strains in which the expression of *STT4* could be repressed with the addition of doxycycline (14). In the presence of this repressor, there was approximately a 10-fold decrease in *STT4* transcript levels and budding growth was similar to that of the wild-type. When *STT4* was repressed, this mutant was able to undergo invasive filamentous growth in response to serum, yet invasive filaments emanating from mutant colonies were ∼ 4-fold shorter compared to wild-type and complemented strains. Upon repression of *STT4* in liquid media containing serum, the cells elongated with protrusions that were roughly ⅓ the length of wild-type cells after 2 hours at 37°C. However, in these repression conditions, the *stt4* mutant still expressed *STT4* and plasma membrane PI(4)P was still detected (14). We also generated a ‘decreased abundance by mRNA perturbation’ (DAmP) allele (23) in *C. albicans* (24), by constructing strains in which one copy of *STT4* was deleted and a dominant nourseothricin resistance marker (SAT1) was integrated just 3’ of the *STT4* stop codon. These DAmP mutants had between 2-4-fold reduction in *STT4* transcript levels compared to a wild-type strain, yet filamentous growth was indistinguishable from the wild-type in liquid and on solid media containing fetal calf serum (Figure S1). More recently, a *stt4* deletion mutant was isolated in a screen for mutants exhibiting hypersensitivity to Hsp90 inhibition *via* geldanamycin, with aberrant filamentation in the presence of geldanamycin or RPMI (17).

To determine if plasma membrane PI(4)P is essential, we attempted to delete this PI-4-kinase, as well as the two other putative components of the Stt4 complex, Efr3 and Ypp1. To generate a *stt4* deletion mutant, we removed the remaining *STT4* copy in the DAmP mutant, by taking advantage of the 3’ SAT1 marker to target homologous recombination. Southern blotting, as well as PCR of genomic DNA (gDNA), confirmed the absence of *STT4* in this homozygous deletion strain (Figure S2A, S2B). In addition, we were able to generate homozygous deletion mutants of *EFR3* and *YPP1*, which were verified by PCR (Figure S3A). RT-PCR revealed the complete absence of *STT4* mRNA transcript, as well as that of *EFR3* and *YPP1* in the respective mutants (Figure S2C, S3B). Interestingly, the levels of mRNA transcripts of other PI- and PIP-kinases and phosphatase, including *PIK1*, *MSS4* and *SAC1* were unaffected in the *stt4* deletion mutant (Figure S2C). These strains all grew with doubling times that were indistinguishable from that of the wild-type strain (81 ± 6 min for the wild-type strain compared to 88 ± 7 min for *stt4Δ/stt4Δ*; 86 ± 5 min for *stt4Δ*/*stt4Δ* + *STT4*; 86 ± 2 min for *efr3Δ/efr3Δ*; 89 ± 1 min for *ypp1Δ/ypp1Δ*), indicating that the PI-4-kinase complex is not necessary for viability or yeast-phase growth in *C. albicans*.

In the presence of serum, however, we observed a striking filamentous growth defect (Figures 1 and 2) in the *efr3*, *ypp1* and *stt4* deletion mutants. The mutants appeared to form short germ tubes but were unable to form longer hyphal filaments. Similarly, all three strains were completely defective in invasive growth agar media containing fetal calf serum (Figure 1C and 2C). These defects were complemented by the reintroduction of a copy of the respective genes, which was confirmed by PCR of gDNA and RT-PCR of mRNA transcripts (Figure S2B, S2C, S3A, S3B). A *stt4* allele that is a hypomorph with reduced *in vivo* lipid kinase activity (25) was identified in *S. cerevisiae* in a genetic screen for aminophospholipid transport mutants (7). Hence, we also examined whether such a mutant, in which a highly conserved amino acid in the catalytic domain, Gly 1782 was changed to Asp (in *C. albicans* G1810D), was critical for hyphal growth. Figure 1B shows that a strain expressing as a sole copy this mutated version of *STT4* exhibited defects in filamentous growth that were intermediate between the *stt4* deletion mutant and the complemented strain. Furthermore, compared to the wild-type and complemented strains, the *efr3*, *ypp1* and *stt4* deletion mutants did not grow in the presence of the cell wall perturbants, including the antifungal drug caspofungin, calcofluor white and Congo Red (Figure 3). A lower concentrations of caspofungin (50 ng/mL compared to 125 ng/mL) revealed that the *stt4* deletion mutant exhibits the greatest sensitivity to this antifungal drug, with the *ypp1* deletion mutant exhibiting intermediate sensitivity and *efr3* deletion mutant having the lowest sensitivity (Figure S4A). These mutants were however not temperature sensitive and grew similar to wild-type controls at 30°C and 37°C (Figure S4B) and, in contrast to what has been previously reported (26), they did not exhibit an increased sensitivity to fluconazole. Given the effects of cell wall perturbants, we examined the cell wall thickness and composition in the *stt4* homozygous deletion mutant. Figures 4A and 4B show that, in this PI-4-kinase deletion mutant, the cell wall was on average 50% thicker compared to wild-type cells. Flow cytometry was used to quantitate the exposed β(1,3)-glucan, as well as the total chitin, mannan and glucan. A significant increase in exposed β(1, 3)-glucan, mannan and glucan content was observed in the *stt4* deletion mutant compared to the wild-type strain (Figure 4C). On average *stt4* mutant cells had a 30% increase in exposed β(1,3)-glucan and a 60% in mannan and glucan content. This cell wall integrity defect and increase in cell surface exposure of β(1,3)-glucan are reminiscent of that observed in the phosphatidylserine synthase mutant *cho1Δ/cho1Δ* (27–30). Together, our results show that the PI-4-kinase complex is critical for filamentous growth and cell wall integrity in *C. albicans*, specifically masking cell wall β(1,3)-glucan.

**Figure 1.**
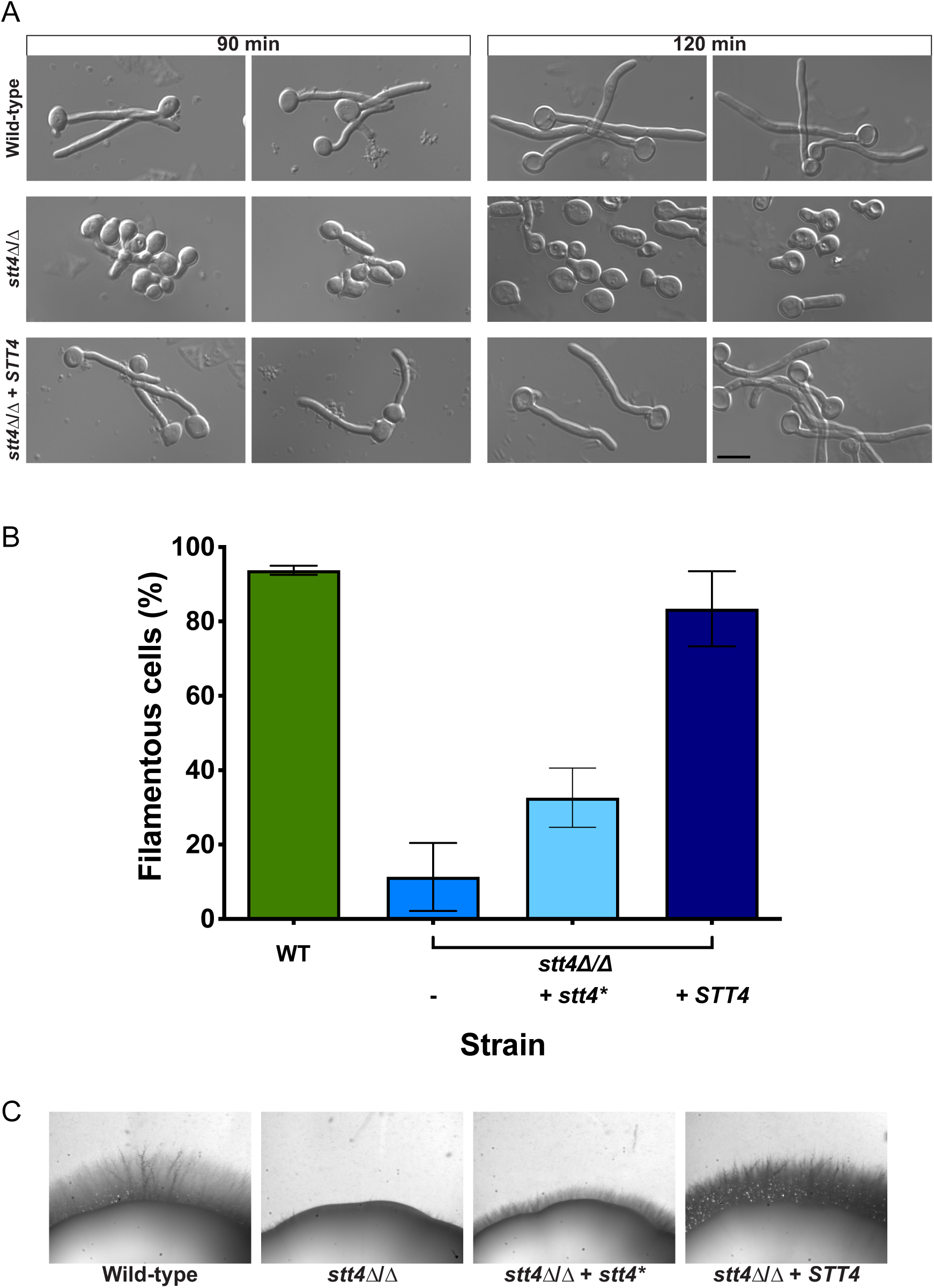
The PI-4-kinase Stt4 is required for filamentous growth. A) Indicated strains (wild-type, PY4861; *stt4*Δ/Δ, PY5111; *stt4*Δ/Δ + *STT4*, PY5131) were incubated with serum at 37°C for 90 and 120 min. B) Percentage of filamentous cells were determined from three independent experiments. (*n* ≥ 120 in each) with strains indicated above in addition to *stt4*Δ/Δ + *stt4\** encoding Stt4[G1810D], PY5757. Cells were considered filametous if germ tubes were twice length of the mother cell or longer, error bars indicate SD. C) Stt4 is required for invasive filamentous growth. Indicated strains were incubated for 4 days at 30°C on serum agar plates. Similar results were observed in three independent experiments.

**Figure 2.**
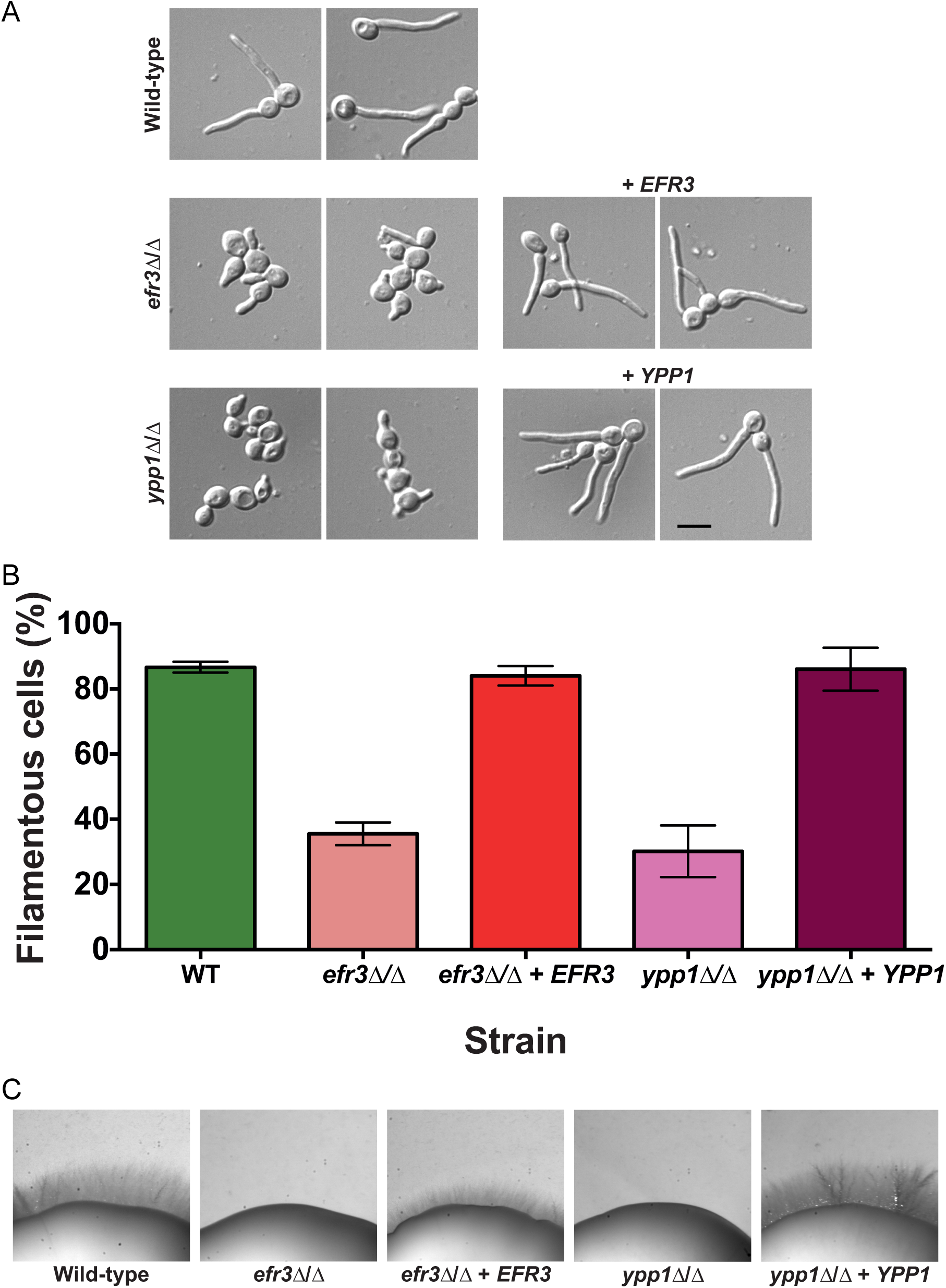
Efr3 and Ypp1 are required for filamentous growth. A) Indicated strains (wild-type, PY4861; *efr3*Δ/Δ, PY4036; *efr3*Δ/Δ + *EFR3*, PY4039; *ypp1*Δ/Δ, PY4033; *ypp1*Δ/Δ +*YPP1*, PY4040) were induced with serum at 37°C for 90 min. B) Percentage of filamentous cells was determined from three independent experiments (*n* ≥ 120 in each). Error bars indicate SD. C) Efr3 and Ypp1 are required for invasive filamentous growth. Indicated strains were incubated for 4 days at 30°C on serum agar plates. Similar results were observed in three independent experiments.

**Figure 3.**
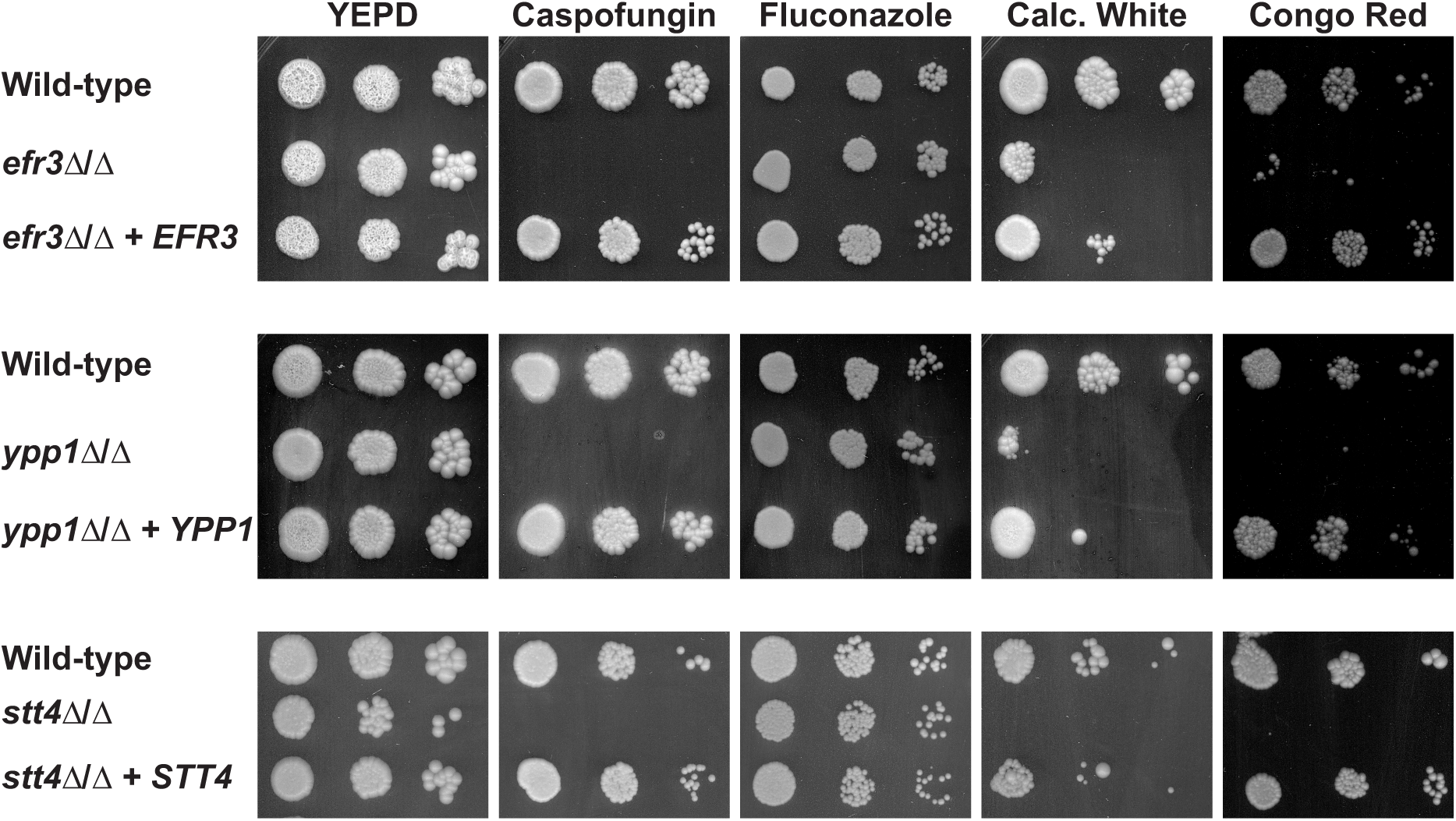
Plasma membrane PI(4)P is important for cell wall integrity. Indicated strains (wild-type, PY4861; *efr3*Δ/Δ, PY4036; *efr3*Δ/Δ + *EFR3*, PY4039; *ypp1*Δ/Δ, PY4033; *ypp1*Δ/Δ +*YPP1*, PY4040; *stt4*Δ/Δ, PY5111; *stt4*Δ/Δ + *STT4*, PY5131) were incubated on YEPD with or without caspofungin (125 ng/mL), fluconazole (10 µg/mL), Calcofluor White (25 µg/mL) or Congo Red (400 µg/mL) for 3 days at 30°C. Similar results were observed in two independent experiments.

**Figure 4.**
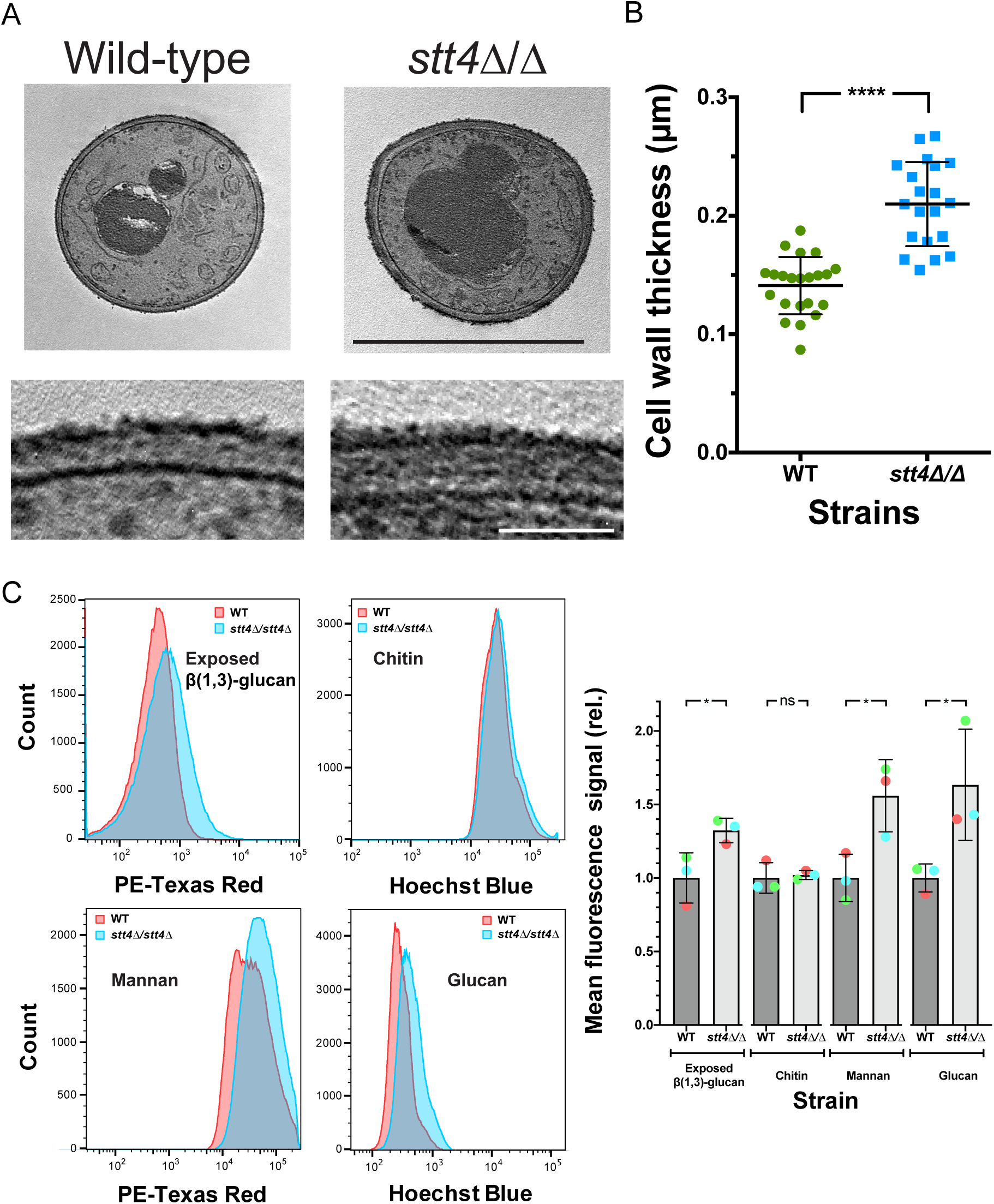
*Stt4* deletion mutant has a thicker cell wall with increased mannan, glucan and exposed β(1,3)-glucan. A) Transmission electron micrographs of the indicated strains (wild-type, PY4861; *stt4*Δ/Δ, PY5111) (upper) with zoom in on the cell wall. Scale bar is 5 µm (upper panel) and 1 µm (lower panel). B) Quantitation of cell wall thickness from EM micrographs. C) The s*tt4* mutant has increased exposure of β(1,3)-glucan together with increased levels of mannan and glucan. Flow cytometry analyses of indicated cells (wild-type, PY4861; *stt4*Δ/Δ, PY5111) labeled with anti-β(1,3)-glucan antibodies and a fluorescently labeled secondary antibody, calcofluor white, fluorescently labeled concanavalin A and aniline blue. Flow cytometry profiles from one biological replicate (10^5^ gated events; left) and means from three biological replicates normalized to each wild-type mean, respectively (right). Bars indicate standard deviations, * is p < 0.05, **** is *p* < 0.0001 and ns is not significant.

### The Stt4 PI-4-kinase complex is specifically required for plasma membrane PI(4)P

The defect in filamentous growth in mutants for all three components of the PI-4-kinase complex suggested that plasma membrane PI(4)P is critical for this process. Therefore, we examined the *in vivo* PI(4)P levels using a fluorescent reporter that binds preferentially this acidic phospholipid at the plasma membrane (13). This reporter can also bind PI(4)P at the Golgi and we have previously shown that, upon a reduction in plasma membrane PI(4)P, the reporter relocalizes to the Golgi (13). In wild-type cells, we observed this GFP-PH^OSH2^-PH^OSH2^-GFP reporter localize predominantly at the plasma membrane, yet in *efr3*, *ypp1* and *stt4* mutants there was a decrease in plasma membrane PI(4)P, with a concomitant increase in intracellular Golgi PI(4)P signal (Figure 5A). Quantification of the reporter fluorescence from these central z-sections, using the Matlab program Hyphal-Polarity (14), indicate that the ratio of mean plasma membrane signal to mean internal signal decreased progressively in the *efr3, ypp1* and *stt4* mutants, resulting from the decrease in plasma membrane PI(4)P together with the overall increase in internal PI(4)P levels (Figure 5B). Closer examination of *stt4* mutant cells revealed little to no plasma membrane PI(4)P (Figure 5A) and this was confirmed by quantification of the normalized fraction of total signal at the plasma membrane, *i.e.* the plasma membrane signal excluding the Golgi cisternae divided by the total cell signal excluding the Golgi cisternae (Figure 5C). The average normalized plasma membrane PI(4)P in *stt4* mutants was very close to 0 (0.078 ± 0.016), with 60% of cells (*n* = 200) having undetectable plasma membrane PI(4)P. Together, these results indicate that *C. albicans* cells with undetectable plasma membrane PI(4)P are viable and suggest that this plasma membrane acidic phospholipid is specifically critical for filamentous growth.

**Figure 5.**
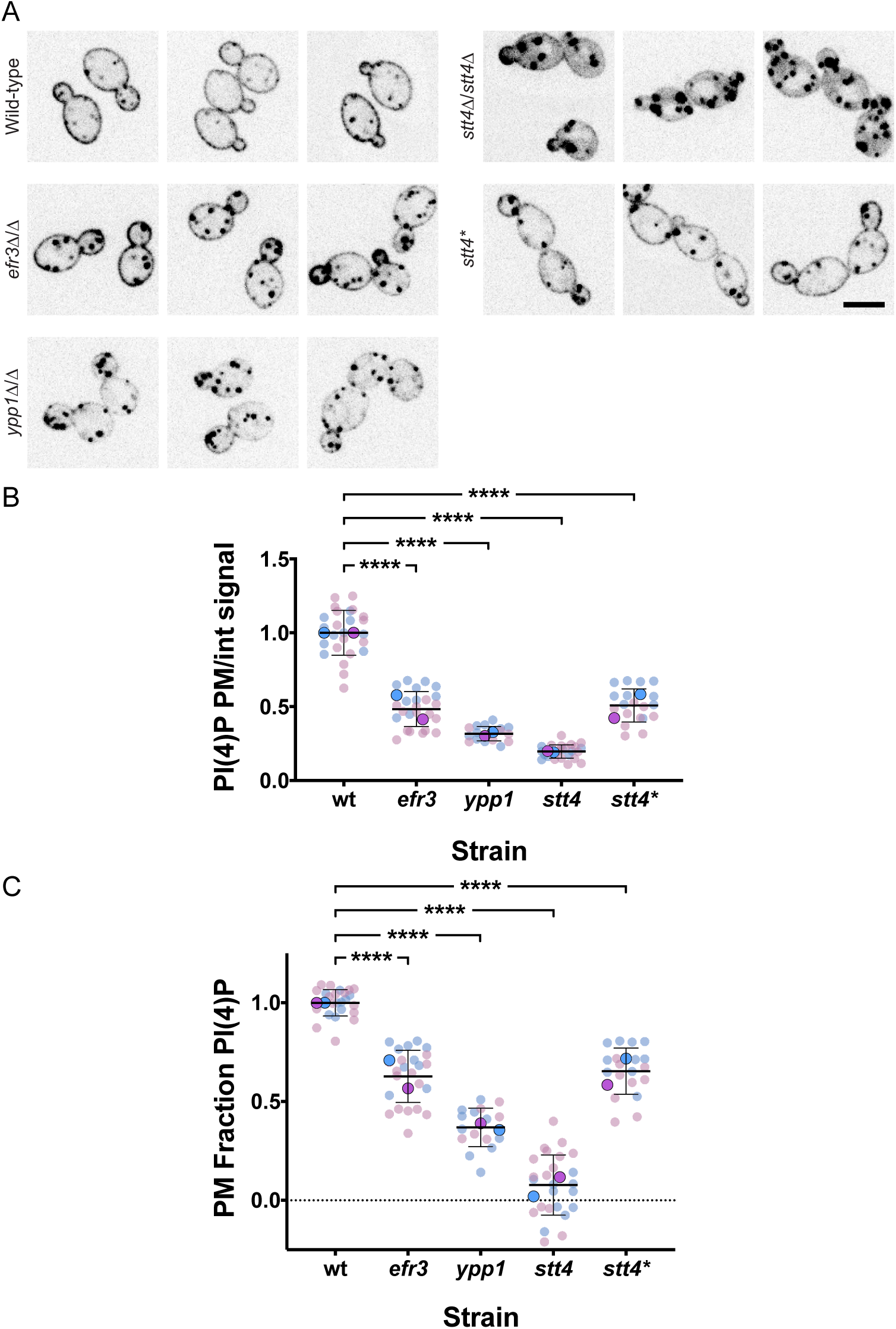
Efr3, Ypp1 and Stt4 are critical for plasma membrane PI(4)P. A) Indicated strains expressing the plasma membrane PI(4)P reporter, GFP-Osh2^PH^-Osh2^PH^-GFP (wild-type, PY2626; *efr3*Δ/Δ, PY4947; *ypp1*Δ/Δ, PY3950; *stt4*Δ/Δ, PY5169; *stt4*Δ/Δ + *stt4\**, PY5838) were imaged and central z-sections of representative cells are shown with inverted look up table (LUT). B – C) Quantitation of plasma membrane and internal signals reveal little to no plasma membrane PI(4)P the *stt4* mutant. The ratio of plasma membrane to internal signal (normalized to the wild-type) and the relative plasma membrane signal (normalized plasma membrane/total signal) is shown. Quantitation of plasma membrane and internal signals was carried out excluding Golgi cisternae. For the wild-type, the mean ratio plasma membrane to internal signal was 3.8 and the ratio plasma membrane divided by total signal was 0.8. We are able to detect ∼1.5% of wild-type plasma membrane PI(4)P levels. Smaller symbols are values from two experiments (6-15 cells per experiment), larger symbols are mean of each experiment with bars indicating overall means and standard deviations. **** is *p* < 0.0001

Previously we have shown that a reduction in Golgi PI(4)P results in Golgi proliferation (13). Hence, we examined whether the reduction of plasma membrane PI(4)P observed in *efr3*, *ypp1* and *stt4* deletion mutants affected the Golgi. While there was a small decrease (20-25%) in the number of Golgi cisternae per cell in *efr3* and *ypp1* mutants compared to the wild-type (Figure 6A), there was no difference in the number of Golgi cisternae per cell in the *stt4* mutant (Figure 6B). These results further confirm that Golgi and plasma membrane PI(4)P pools are functionally distinct. Our previous analyses of a *STT4* repressible strain revealed that upon a 10-fold repression of *STT4* transcript, there was a reduction of plasma membrane PI(4,5)P_2_ (14). As a result, we examined plasma membrane PI(4,5)P_2_ levels in the *efr3*, *ypp1* and *stt4* deletion mutants. Figure 7 shows that this phosphatidylinositol phosphate was observed at the plasma membrane in all three mutants. Compared to the wild-type strain, there was a reduction of 10-15% in the plasma membrane PI(4,5)P_2_ levels to internal ratios in the efr3, *ypp1* and *stt4* mutants. Comparison of the fraction of plasma membrane PI(4,5)P_2_ signal revealed a small (10% or less) reduction in the *stt4* mutants with respect to the wild-type strain, similar to our previous observations with a repressible strain (14). Together these results demonstrate that the dramatic reduction of plasma membrane PI(4)P does not alter Golgi PI(4)P nor does it substantially alter plasma membrane PI(4,5)P_2_.

**Figure 6.**
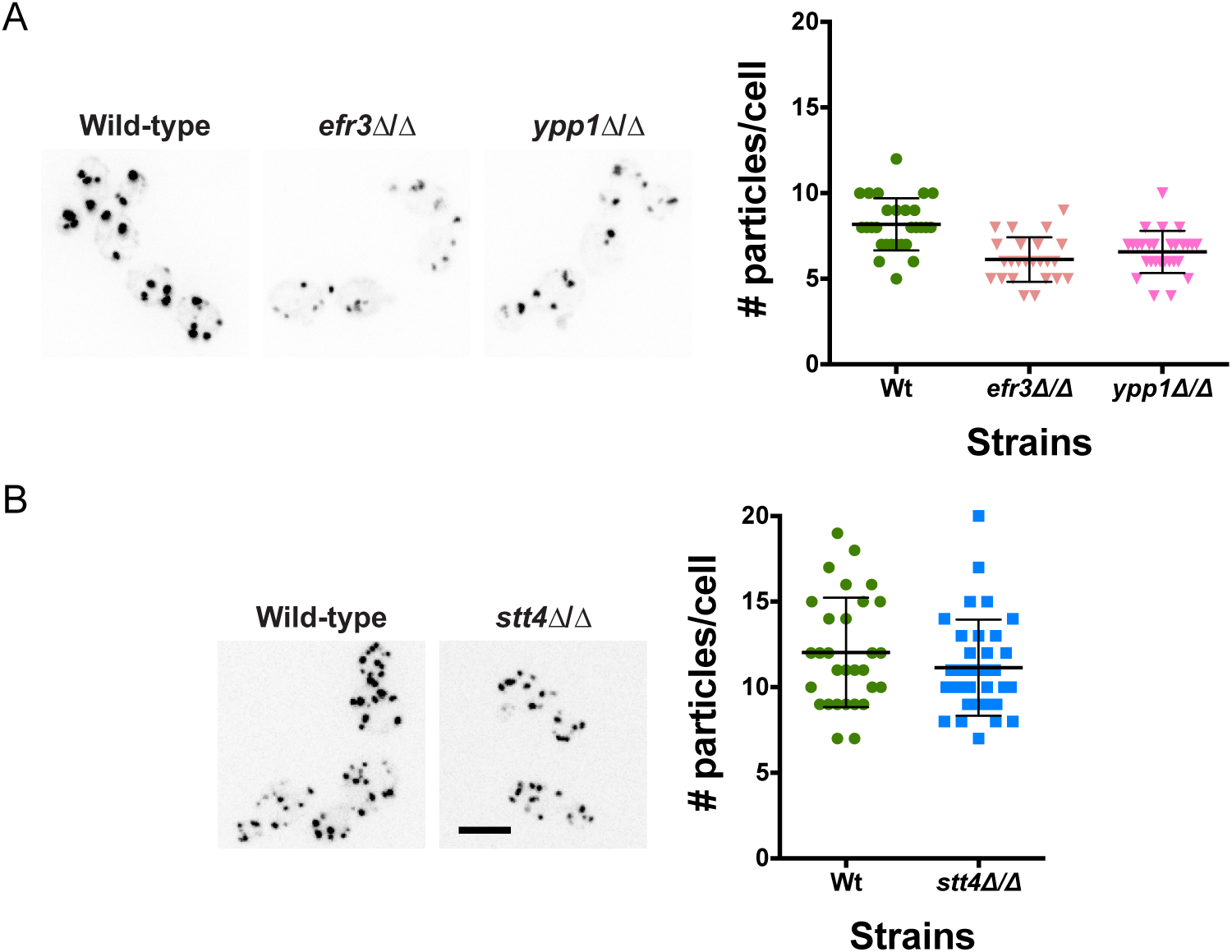
The number of Golgi cisternae is not affected by a decrease in plasma membrane PI(4)P. A-B) Indicated strains expressing Golgi PI(4)P reporter, FAPP1-GFP (wild-type, PY2578; *efr3*Δ/Δ, PY3933; *ypp1*Δ/Δ, PY3951; *stt4*Δ/Δ, PY5552) were imaged and maximum projections of representative cells are shown with inverted LUT (left). Quantitation of the number of Golgi cisternae per cell (right) in the indicated strains (*n* = 24 -34 cells per strain).

**Figure 7.**
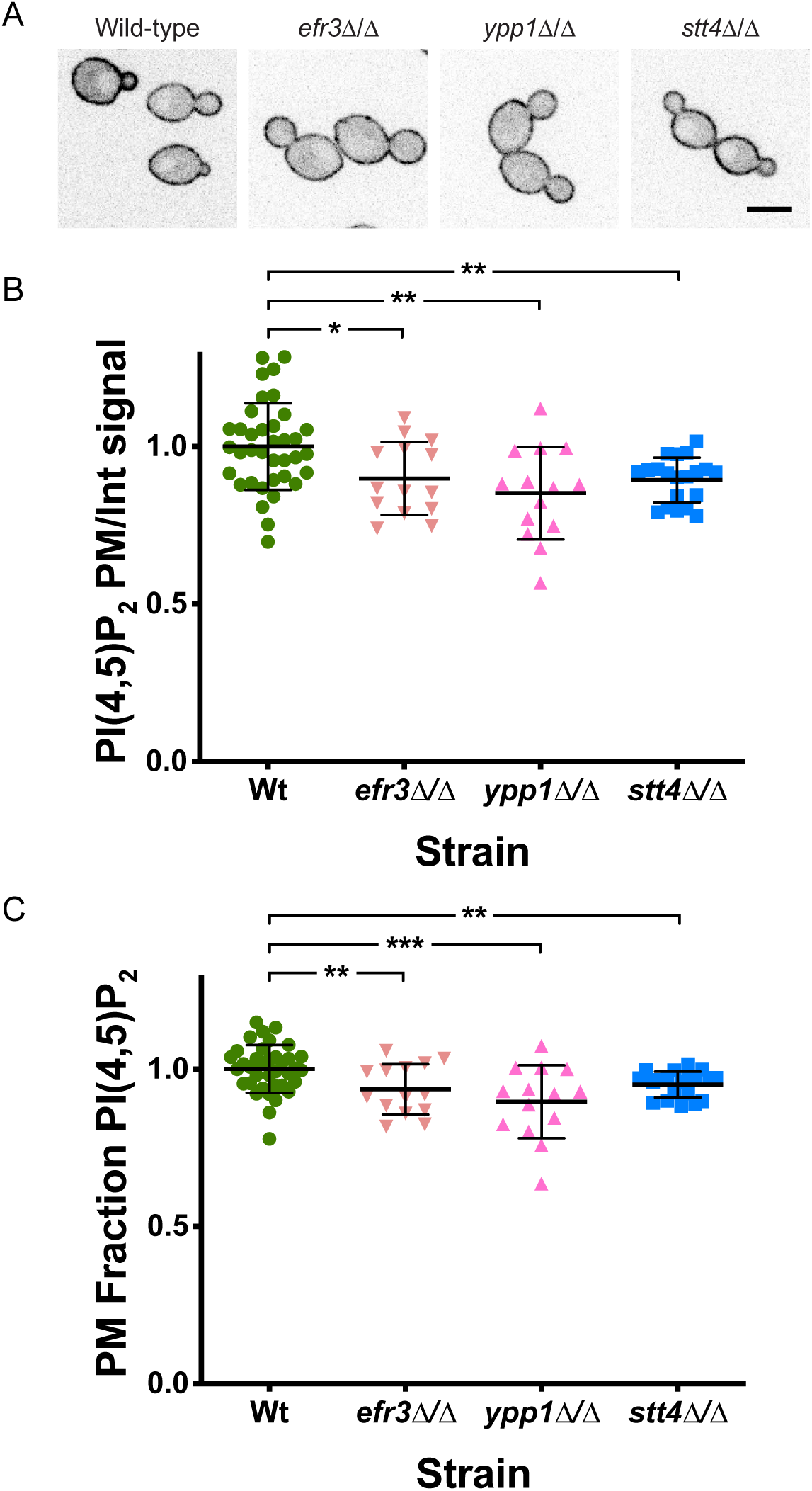
Plasma membrane PI(4,5)P_2_ is not substantially affected by a decrease in PI(4)P. A) Indicated strains expressing PI(4,5)P_2_ reporter, GFP-PH^Plcδ^-PH^Plcδ^-GFP wild-type, PY1206; *efr3*Δ/Δ, PY3935; *ypp1*Δ/Δ, PY3958; *stt4*Δ/Δ, PY555) were imaged and central z-sections of representative cells are shown with inverted LUT. B–C) Quantitation of plasma membrane and internal signals reveals that plasma membrane PI(4,5)P_2_ is largely unaffected in the absence of Efr3, Ypp1 and Stt4. The ratio of plasma membrane to internal signal and the relative plasma membrane signal determined as in Figure 5B, 5C (*n* = 15-20 cells; 2 experiments for WT). For the wild-type, the mean ratio plasma membrane to internal signal was 2.8 and the ratio plasma membrane divided by total signal was 0.7. * < 0.02, ** is p < 0.01 and *** is *p* < 0.0005.

Oxysterol binding proteins, such as Osh6 and Osh7 in *S. cerevisiae*, are lipid transfer proteins that transfer phosphatidylserine (PS) from the ER to the plasma membrane concomitant with transfer of PI(4)P from the plasma membrane to the ER, where it is hydrolyzed by Sac1 (31–33). In *C. albicans*, as in *S. cerevisiae*, Sac1, which localizes to the ER and Golgi, is critical for regulating plasma membrane PI(4)P levels (13, 34, 35). Given that in the *efr3*, *ypp1* and *stt4* deletion mutants there was a reduction in PI(4)P levels, we examined whether PS plasma membrane levels were affected using a fluorescent reporter that binds preferentially this acidic phospholipid (36). In wild-type cells we observed this LactC2-GFP reporter localized predominantly at the plasma membrane with little to no internal signal observed (Figure 8A). In contrast, in the *efr3*, *ypp1* and *stt4* mutants, peri-nuclear signal was observed, characteristic of the ER, yet plasma membrane PS was still apparent in each of these mutants (Figure 8A). Quantification of signals from central z-sections using the Matlab program Hyphal-Polarity (14) confirmed that the ratio of mean plasma membrane signal to mean internal signal decreased progressively in the *efr3, ypp1* and *stt4* mutants, resulting in part, from a progressive increase in internal PS signal (Figure 8B). Similarly, the mean plasma membrane PS fraction also decreased progressively in the *efr3, ypp1* and *stt4* mutants, with the *stt4* deletion mutant exhibiting an approximately 50% reduction compared to wild-type cells. The *stt4* hypomorph, Stt4[G1810D], had plasma membrane PS levels intermediate between the *efr3* and *ypp1* mutants. Together, these results suggest that sufficient PI(4)P is critical for the transport of PS from the ER to the plasma membrane. We next examined if there was a correlation between the plasma membrane PI(4)P and PS levels. Figure 9 shows that there is a direct correlation between the levels of these two lipids in the different deletion mutants and the wild-type strain. Plasma membrane lipid levels in the Stt4[G1810D] mutant were also consistent with such a correlation. Note that while we were unable to detect plasma membrane PI(4)P in the *stt4* deletion strain, approximately 50% of the plasma membrane PS was detectable in this mutant. Furthermore, plasma membrane PS levels in the *efr3* mutant were not dramatically different from that of the wild-type cells. Together, our results suggest that the filamentation and cell wall integrity defects observed in the three Stt4 complex mutants are likely to be due to lack of plasma membrane PI(4)P and not PS.

**Figure 8.**
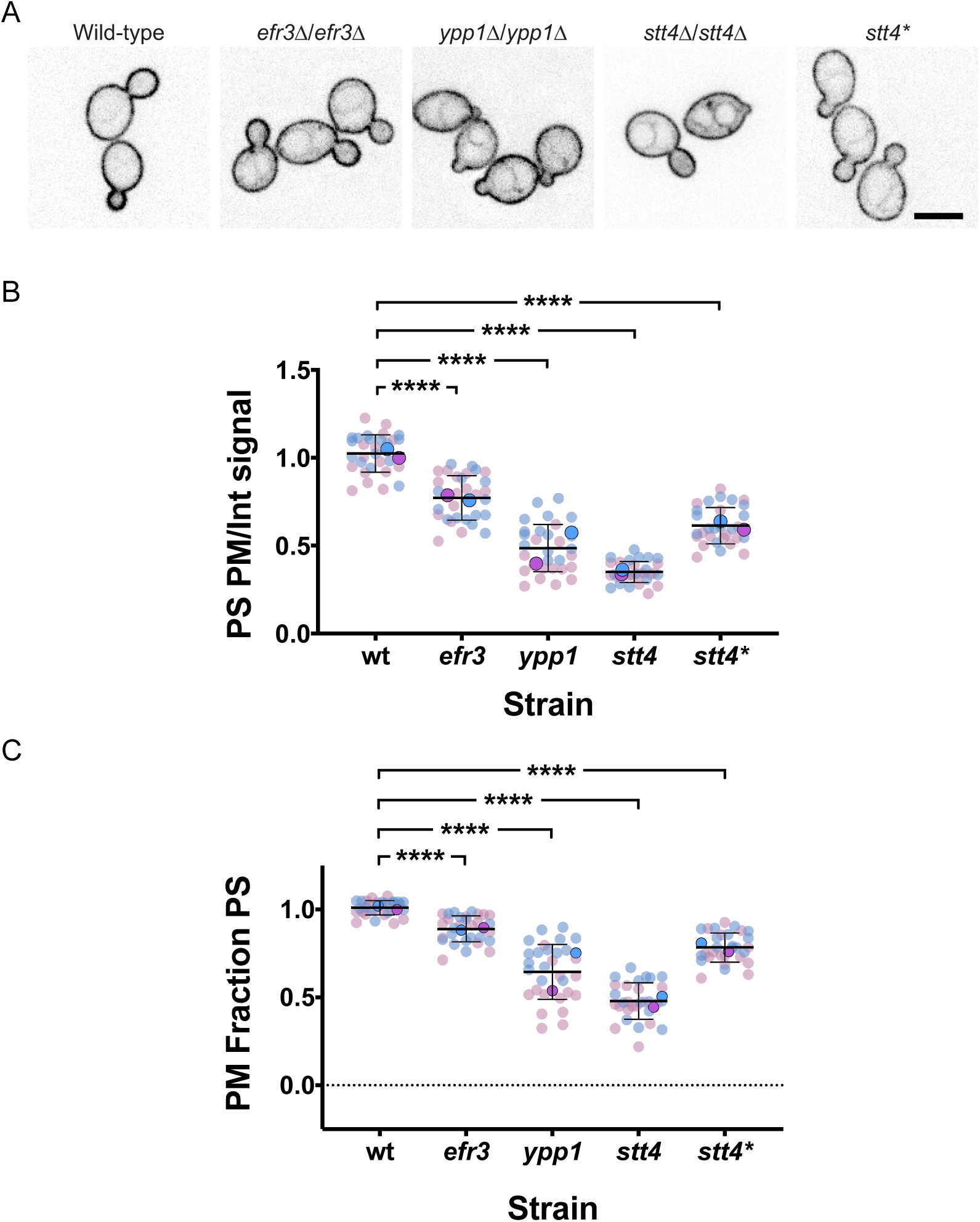
A reduction in plasma membrane PI(4)P results in an increase in PS at the ER. A) Indicated strains expressing PS reporter GFP-LactC2 (wild-type, PY3239; *efr3*Δ/Δ, PY4124; *ypp1*Δ/Δ, PY4131; *stt4*Δ/Δ, PY5174; *stt4*Δ/Δ + *stt4\**, PY5903) were imaged and central z-sections of representative cells are shown with inverted LUT. B-C) Quantitation of plasma membrane and internal signals reveals a progressive decrease in plasma membrane PS in *efr3*, *ypp1* and *stt4* strains. The ratio of plasma membrane to internal signal and the relative plasma membrane signal determined as in Figure 5B, 5C. For the wild-type, the mean ratio plasma membrane to internal signal was 4.5 and the ratio plasma membrane divided by total signal was 0.8. Smaller symbols are values from two experiments (*n* = 15 cells each), larger symbols are mean of each experiment with bars indicating overall means and standard deviations.**** is *p* < 0.0001

**Figure 9.**
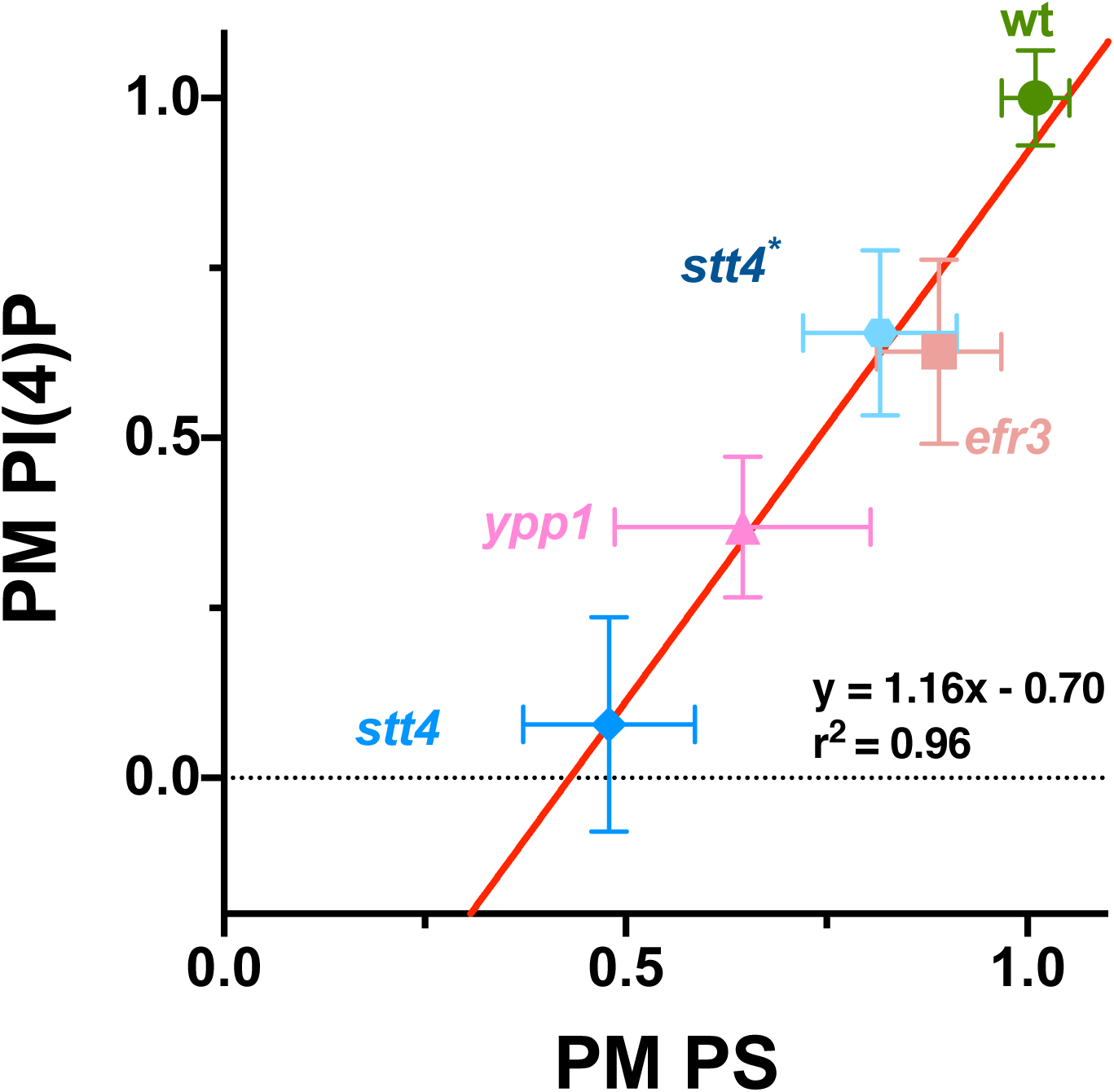
Plasma membrane PS is proportional to PI(4)P levels. Average levels of plasma membrane PS and PI(4)P in indicated strains were normalized to 1 in the wild-type. Linear curve fit: *y* = 1.316*x* − 0.70; r^2^ = 0.96. Bars are standard deviation with *n* = 16 - 40 cells for each determination.

### Stt4 PI-4-kinase complex localizes to cortical patches

To determine the distribution of Efr3, Ypp1 and Stt4, we generated 3x-mScarlet fusions by tagging the chromosomal copy of the respective genes. These fusion were functional in that as a sole copy they complemented the cell wall integrity defect of the respective mutants (Figure S5). Despite the low abundance of these Stt4 PI-4-kinase complex subunits, we observed patches around the cortex of the mother cell and buds (Figure 10A), which were also visible along the cortex of the germ tubes with reduced signals in the mother cell (Figure 10B). In *S. cerevisiae,* Ypp1 and Efr3 are critical for Stt4 membrane localization. Here, we observed that while Stt4 localization to the cortex is dependent on Ypp1 in *C. albicans*, in the *efr3* mutant there were still some cells with cortex localized Stt4 (Figure 10C). Efr3 cortex localization depended upon Ypp1 and Stt4, with loss of cortex signal observed in either mutant. In contrast, there were some cells with Ypp1 cortex signal in the *efr3* mutant, but not in the *stt4* mutant. In this *efr3* mutant 40-50% of cells exhibited punctate localization of Stt4 and Ypp1. In the *ypp1* mutant, we did not observe cells with either Stt4 or Efr3 localized. Finally, in the *stt4* mutant, only ∼10% of cells exhibited punctate localization of Ypp1. RT-PCR revealed that the mScarlet transcript levels in all mutants were similar to that of the wild-type (Figure S6). As these results suggested that Ypp1 and Stt4 were the most critical components of this PI-4-kinase complex, we examined whether these proteins colocalized to the same cortical patches. Figure S7 shows that, in a strain expressing Ypp1-mTurquoise and Stt4-3xmScarlet, there was only limited colocalization of these two proteins, suggesting that, if they form a complex, it is likely to be transient.

**Figure 10.**
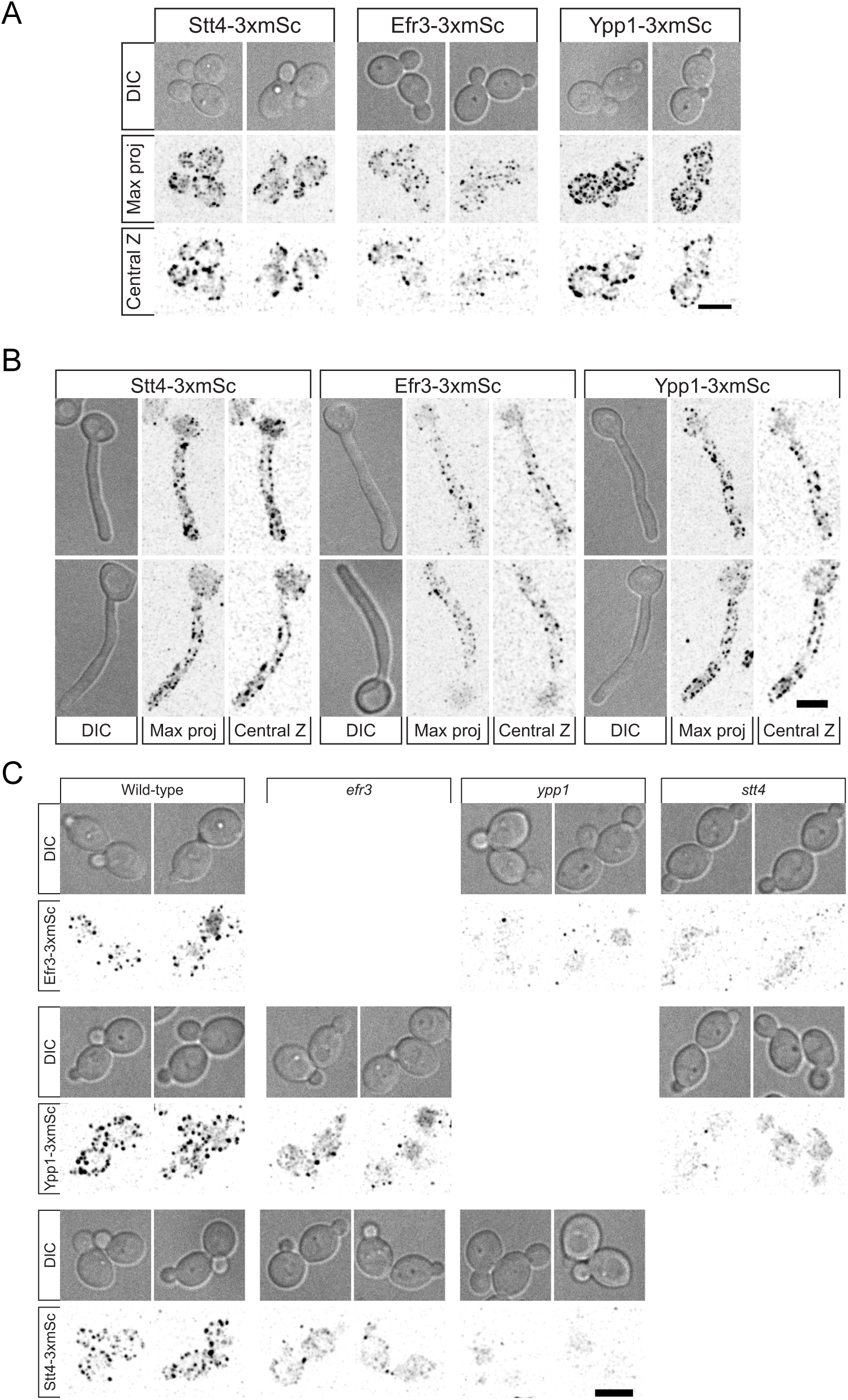
Efr3, Ypp1 and Stt4 localize as cortical patches, with Ypp1 and Stt4 critical for each other’s localization. A-B) Strains expressing indicated 3xmScarlet fusions (Stt4-3xmSc, PY6193; Efr3-3xmSc, PY6197; Ypp1-3xmSc, PY6195) were imaged during budding (A) and hyphal (B) growth. DIC images, central z-sections and maximum projections of 17 x 0.5 µm z-sections are shown. C) Indicated strains (WT, PY6197, PY6195, PY6193; *efr3*, PY6136, PY6142; *ypp1*, PY6138, PY6144; *stt4*, PY6140, PY6134) strains expressing respective 3xmScarlet fusions were imaged during budding growth and maximum projections of 17 x 0.5 µm z-sections are shown.

### Plasma membrane PI(4)P is critical for virulence

To investigate the importance of plasma membrane PI(4)P in virulence, we examined the *efr3, ypp1* and *stt4* mutants in two murine infection models, HDC and oropharyngeal candidiasis (OPC). As *C. albicans* responds to cues, such as the presence of serum, by the induction of hyphal specific genes (HSG), many of which are critical for virulence, we initially examined HSG levels after 30 and 120 min incubation with serum. Figure 11A shows that the *stt4* deletion mutant induced a range of HSGs, including *ECE1*, *HGC1*, *HWP1*, *ALS3* and *SAP4-6*, 10^3^ – 10^5^-fold (excluding *SAP4-6*), similar to the wild-type strain, with only a 5-fold reduction in induced transcript levels observed after 30 min incubation. In the HDC model, 20% and 100% of mice infected with the *ypp1* and *stt4* mutants, respectively, survived 2 weeks after injection, while all of the mice infected with the *efr3* mutant or wild-type strain died within 5-6 days (Figure 11B). The virulence was significantly restored in complemented strains (Figure 11B). Furthermore, we examined whether the *stt4* deletion mutant could filament after long incubation times in serum or in kidney homogenate (37). After 6 Hr incubation with either serum or kidney homogenate, despite observing elongated cells, the *stt4* mutant did not form hyphal filaments, compared to wild-type and complemented strains (Figure S8). It is unlikely that the unmasking of cell surface β(1,3)-glucan and these virulence defects are due to altered plasma membrane PS levels as a *cho1Δ/cho1Δ::CHO1* strain with greater than 50% reduction in PS levels had no increase in unmasked β(1,3)-glucan and exhibited full virulence in mouse models of systemic and oropharyngeal infection (27–29). In the OPC model, only the *ypp1* mutant was substantially less virulent than the wild-type strain, with the oral fungal burden of the infected mice reduced by ∼25-fold (Figure 11C). Nonetheless, examination of the histopathology of tongue thin sections revealed that in infection lesions, all strains were able to filament (Figure S9). Together, these data suggest that plasma membrane PI(4)P is required for virulence during hematogenously disseminated candidiasis, with a decrease in virulence observed upon a 60% reduction in plasma membrane PI(4)P and the lack of lethality observed in the absence of this phosphatidylinositol phosphate. Our data also indicate that only a small amount of plasma membrane PI(4)P is required for normal virulence during oropharyngeal candidiasis.

**Figure 11.**
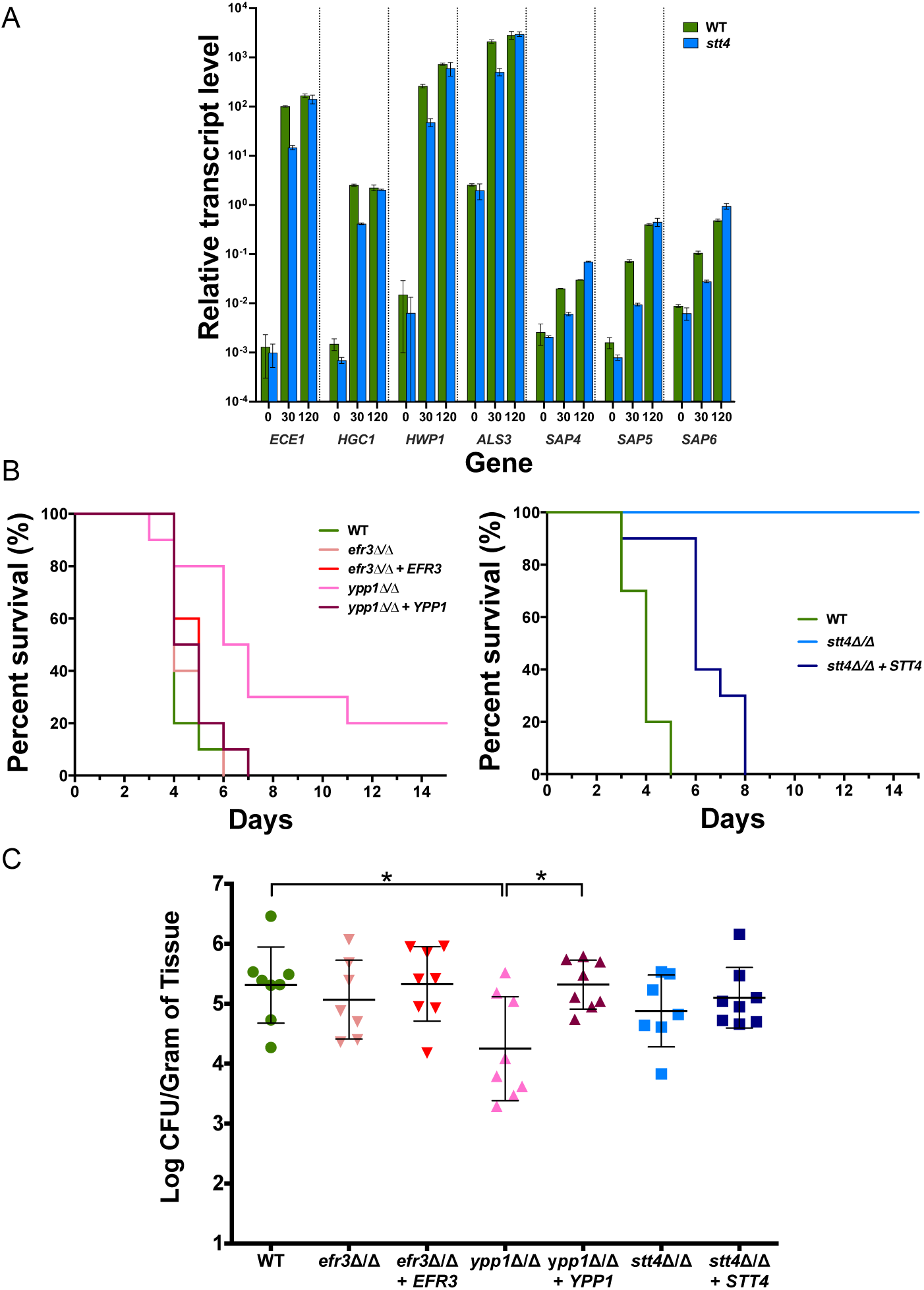
Plasma membrane PI(4)P is specifically required for hematogenously disseminated candidiasis. A) Hyphal specific genes are induced in a *stt4* deletion mutant. The transcript level of indicated hyphal specific genes was determined in wild-type (PY4861) and *stt4*Δ/Δ (PY5111) strains at indicated times (min) with serum at 37°C by qRT-PCR and normalized to the levels of *TDH3* transcript. Means of 3 determinations from an experiment are shown with bars indicating standard deviations. Similar results were observed in two additional biological replicates. B) Stt4 is required for virulence in a murine systemic infection model. Survival of mice (*n* = 10) over time following injection of indicated strains (wild-type, PY4861; *efr3*Δ/Δ, PY4036; *efr3*Δ/Δ + *EFR3*, PY4039; *ypp1*Δ/Δ, PY4033; *ypp1*Δ/Δ +*YPP1*, PY4040; *stt4*Δ/Δ, PY5111; *stt4*Δ/Δ + *STT4*, PY5131). Similar results were observed in two independent experiments, differences between WT and *ypp1Δ*/*Δ* was statistically significant, *p* = 0.002 (left panel) and between WT and *stt4Δ*/*Δ* was statistically significant, *p* < 0.0001 (right panel). C) Stt4 and Efr3 are not required for virulence in a murine oropharyngeal infection model. Colony forming units per gram of tongue tissue, 5 days subsequent to oropharyngeal infection (*n* = 8 mice per strain) with indicated strains (wild-type, PY4861; *efr3*Δ/Δ, PY4036; *efr3*Δ/Δ + *EFR3*, PY4039; *ypp1*Δ/Δ, PY4033; *ypp1*Δ/Δ +*YPP1*, PY4040; *stt4*Δ/Δ, PY4414; *stt4*Δ/Δ + *STT4*, PY4433). * is *p* < 0.05.

## Discussion

Our results show that plasma membrane PI(4)P is critical for the *C. albicans* yeast to filamentous growth transition and cell wall integrity. We show that all three members of the Stt4 PI-4-kinase complex are dispensable for viability, yet are required for filamentous growth and cell wall integrity. Furthermore, quantitative analyses indicate that these mutants have decreasing levels of plasma membrane PI(4)P, going from *efr3* to *ypp1* to *stt4*; a majority of the *stt4* cells lack detectable PI(4)P at the plasma membrane. In addition, a *stt4* hypomorph (25) had similar plasma membrane PI(4)P levels as the *efr3* mutant. We observed little to no alteration in Golgi PI(4)P and plasma membrane PI(4,5)P_2_ in these Stt4 PI-4-kinase complex mutants. Consistent with a link between plasma membrane PI(4)P and PS, there is a gradual increase in internal pools of PS in *efr3, ypp1* and *stt4* mutants, yet even in the *stt4* mutant that lacks plasma membrane PI(4)P, PS is still detected at the plasma membrane. All three of these PI-4-kinase complex proteins localize to cortical patches but only Ypp1 and Stt4 appear to be critical for the complex formation. Furthermore, our results reveal that plasma membrane PI(4)P is important for masking cell surface β(1,3)-glucan, but not for induction of a number of hyphal specific genes. Plasma membrane PI(4)P is critical for pathogenicity during hematogenously disseminated candidiasis, but less so for oropharyngeal candidiasis, suggesting that this lipid has different roles in distinct anatomic infection sites, which we attribute, in part, to host immune recognition *via* unmasked cell surface β(1,3)-glucan.

Using strains in which either the *C. albicans* PI-4-kinase, *i.e.* Pik1 at the Golgi or Stt4 at the plasma membrane, could be repressed, we previously showed that the Golgi PI-4-kinase is strictly required for invasive filamentous growth (13), whereas repression of the plasma membrane PI-4-kinase mutant resulted in cells that can still form short protrusions and invasive filaments (14). This repressible *stt4* mutant had an ∼10-fold reduction in *STT4* transcript levels compared to a wild-strain, yet PI(4)P was still detectable at the plasma membrane. To address the importance of plasma membrane PI(4)P in *C. albicans,* we generated homozygous deletion mutants in the PI-4-kinase complex, comprised of Efr3, Ypp1 and Stt4. None of these PI-4-kinase components are essential for viability in *C. albicans* in contrast to other fungi, specifically in *S. cerevisiae* where both Pik1 and Stt4 are essential (6, 7, 38, 39). We speculate that the plasma membrane PI(4)P is required to maintain filamentous growth in *C. albicans via* contributions to cell polarity and membrane traffic. *A. nidulans* and *C. neoformans* appear to only have a Stt4 PI-4-kinase, that is essential for viability (4, 5). In addition, although *C. albicans stt4*, *ypp1*, and *efr3* mutants showed increased sensitivity to cell wall perturbants, they did not exhibit a temperature sensitive growth defect, as was the case in *S. cerevisiae* where a temperature sensitive *stt4* mutant lysed at the non-permissive temperature (15). Our quantitative analyses revealed that the majority of *C. albicans* cells lacking Stt4 have undetectable levels of PI(4)P at the plasma membrane, suggesting that this lipid is dispensable for viability. Such *C. albicans* mutants with little to no plasma membrane PI(4)P are sensitive to cell wall perturbants and have thicker cell walls with increased levels of glucan and mannan, along with an increase in cell surface exposed β(1,3)-glucan, indicating that PI(4)P is critical for maintaining cell wall integrity. We speculate that plasma membrane glucan synthase activity is regulated by PI(4)P, *via* the Rho1 regulatory subunit (40). This raises the question of how PI(4,5)P_2_ is generated in the *stt4* deletion mutant and we speculate that PI(4)P from the Golgi may be phosphorylated by Mss4. This could occur *via* Mss4 localization to the Golgi or secretory vesicles, although this kinase has not been detected in these compartments in *C. albicans* (14). Alternatively, a small amount of Golgi PI(4)P could reach the plasma membrane and be immediately phosphorylated.

Previous studies in *S. cerevisiae* indicated that the levels of PS and plasma membrane PI(4)P are linked. For example, an *S. cerevisiae stt4* mutant with reduced levels of plasma membrane PI(4)P accumulates PS (7, 25). Furthermore a *S. cerevisiae sac1* phosphatase mutant that results in a dramatic increase in PI(4)P (34, 35) has decreased PS levels at the plasma membrane, with an increase in intracellular membranes (41). Similarly, in fibroblasts, PI-4-kinase III*α* knockouts have decreased plasma membrane PS (42). The *S. cerevisiae* oxysterol proteins Osh6 and Osh7 have been shown to exchange PS for PI(4)P *in vitro*, and *in vivo* a *sac1* mutation resulted in a redistribution of added lyso-PS, which is normally at the plasma membrane, to the ER (32). It was proposed that a PI(4)P gradient from the plasma membrane to the ER drives PS transport *via* Osh6/7 from the ER to the plasma membrane (32). Given that *C. albicans* cells are viable with little to no plasma membrane PI(4)P, it is likely that this so-called ‘phosphoinositide-motive force’ is not essential (43). Nonetheless, we observed a progressive increase in intracellular PS in *C. albicans* mutants with decreasing plasma membrane PI(4)P, demonstrating the plasma membrane PI(4)P is critical for PS transport to the plasma membrane. This result indicates that Osh6/7 counter-transporters account for roughly half of the plasma membrane PS and suggests the remaining PS may be delivered to the plasma membrane *via* vesicular traffic.

In *S. cerevisiae,* Stt4 and Ypp1 and Efr3 and Ypp1 colocalize in cortical patches (18) and the latter interaction has also been observed by bimolecular fluorescence complementation (44). In mammalian cells and fission yeast, Stt4, Ypp1 and Efr3 also form a complex observed by co-immunoprecipitation and co-localization (8, 19, 45, 46). Here, we show that all three Stt4 PI-4-kinase complex proteins localize to cortical patches during budding and hyphal growth, yet we did not observe substantial co-localization between Ypp1 and Stt4 proteins, although each protein was critical for the cortical patch localization of the other protein. This suggests that Ypp1 is important for targeting and/or stabilization of the Stt4 PI-4-kinase, as has been observed in *S. cerevisiae* (18). Interestingly, although Efr3 is important for cortical patch localization of Ypp1 and Stt4 proteins, it is not absolutely required as in *S. cerevisiae* (18). These results suggest that Efr3 facilitates targeting and/or stabilization of the complex, but that plasma membrane PI-4-kinase activity is not strictly dependent on it.

Cell wall defects in mutants lacking the PS synthase Cho1, that have little to no PS, are in some respects similar to those of *stt4* deletion mutant cells. For example, a *C. albicans cho1* deletion mutant also had a thicker cell wall and exhibited increased sensitivity to the antifungal drug caspofungin (27). However, this *cho1* mutant had a dramatic increase in cell wall chitin levels (27, 30, 47), in contrast to a mutant lacking plasma membrane PI(4)P. Interestingly, both *stt4* and *cho1* deletion mutants exhibited an increase in exposed cell wall β(1,3)-glucan, with the latter mutant having a roughly 10-fold greater increase in this polysaccharide (30) compared to the *stt4* mutant. However, given that a *cho1Δ*/*cho1Δ*::*CHO1* strain with >50% reduction in PS levels had no increase in unmasked β(1,3)-glucan (27, 28), it is unlikely that the cell wall defects observed in the mutant lacking plasma membrane PI(4)P is due to the reduced PS levels.

The importance of lipids with respect to *C. albicans* virulence has been challenging to determine, as a number of lipids are essential or viability and cell growth. One lipid that has been shown to be critical for pathogenicity in a range of different fungi is the sphingolipid glucosylceramide (48–51). Furthermore, mutants lacking either the PS synthase Cho1 or PS decarboxylases (Psd1 and Psd2) were avirulent in murine models of systemic candidiasis and oropharyngeal candidiasis (27, 29). However, both of these mutants grew substantially slower than wild-strains and exhibited a ∼50% reduction in phosphatidylethanolamine (PE) levels (27). Expression of a heterologous serine decarboxylase revealed that the observed virulence defects, as well as β(1,3)-glucan unmasking, were in fact due to the ethanolamine auxotrophy of these mutants (29). Using an *in vitro* assay for host pathogen interactions, an ∼ 2-fold reduction in macrophage lysis was observed with a *stt4* mutant (17), suggesting that this kinase may be important for virulence. Similarly, the filamentation defect and increased exposure of cell surface β(1,3)-glucan in the plasma membrane PI-4-kinase deletion mutant indicated that PI(4)P has a role in virulence. Interestingly, a recent study by Dunker *et al.* showed that rapid proliferation can compensate for the absence of filamentation in a murine model of systemic candidiasis, directly challenging the idea that filamentation is strictly required for virulence (37). Of the three Stt4 PI-4-kinase complex mutants, *stt4,* and to a lesser extent *ypp1,* exhibited virulence defects in a murine model of systemic candidiasis. Given that all Stt4 PI-4-kinase complex mutants are defective in filamentation, this decrease in virulence is unlikely to be attributable to a defect in morphogenesis. Indeed, elongated *stt4* mutant cells were observed after long incubation times with serum or kidney homogenate, and hyphal gene induction was not substantially different from the wild-type strain after two hours incubation in serum. Furthermore, in a murine model for oropharyngeal candidiasis, all Stt4 PI-4-kinase complex mutants were able to filament. Our results, however, indicate a correlation between PI(4)P at the plasma membrane and systemic candidiasis, given the *ypp1* mutant and the *stt4* mutant have a 3-fold or greater decrease in plasma membrane PI(4)P. As the Stt4 PI-4-kinase complex mutants all have altered cell wall integrity, substantial alterations in the level of plasma membrane PI(4)P lead to corresponding changes in the cell wall, in particular, exposure of masked β(1,3)- glucan that is recognized by host immune cells, ultimately contributing to the reduction in virulence in the systemic candidiasis model. Consistent with this explanation, the *stt4* deletion mutant exhibited little to no defect in OPC infection assay, in which the host immune response is less critical. It will be interesting to further analyze the cell wall defects in these Stt4 PI-4-kinase complex mutants given their different levels of plasma membrane PI(4)P, which will be useful tools in dissecting how plasma PI(4)P regulates the cell wall during fungal infection.

## Acknowledgements

We thank J. Wendland, S. Bates and A. Mitchell for strains and plasmids. We thank S. Bogliolo for assistance. We thank the Platforms Resources in Imaging and Scientific Microscopy facility (PRISM; M. Mondin, S. Lachambre, B. Monterroso), the Cytometry Platform (BV-CyAn; A. Loubat), the Experimental Histopathology Platform (S. Rekima) and Microscopy Imaging Côte d’Azur (MICA) for microscopy, cytometry and histopathology support. This work was supported by the CNRS, INSERM, Université Côte d’Azur and ANR (ANR-11-LABX-0028-01, ANR-16-CE13-0010-01 and ANR-19-CE13-0004-01) grants, by grant R01DE026600 from the NIH USA and grant SAF2017-86192 from Spanish Ministry from Science and Innovation.

## Materials and Methods

### Strain and plasmid construction

Standard methods were used for *C. albicans* cell culture, molecular and genetic manipulations. Strains were grown in yeast extract-peptone dextrose (YEPD) at 30°C unless otherwise indicated and induction of filamentous growth was carried out with 50% serum at 37°C for 90 and/or 120 min. To determine doubling times cells were grown in YEPD media at 30°C and optical density was followed over 8 Hr of logarithmic growth. For induction with kidney homogenates, kidneys from male C57BL/6 mice were aseptically removed, dounce homogenized with sterile PBS (0.4 g/mL) and this homogenate was diluted 1:1 with logarithmically growing *C. albicans* strains in YEPD. Serial dilutions of the different strains on YEPD plates containing Congo red (400 μg/mL), Calcofluor White (25 μg/mL), Caspofungin (50 and 125 ng/mL), and Fluconazole (10 μg/mL) were examined after 2-3 days of incubation at 30°C (52). The strains and plasmids used are listed in Tables S1 and S2, respectively and the oligonucleotides used are listed in Table S3. All strains are based on BWP17 background (53). The *efr3Δ/efr3Δ* and *ypp1Δ/ypp1Δ* strains were generated by homologous recombination. Each copy was replaced by either *HIS1* or *URA3* using knockout cassettes generated by amplification from pGemHIS1 and pGemURA3 (53) with primers CaEfr3pKO/CaEfr3mKO and CaYpp1pKO/CaYpp1mKO. In order to generate prototrophic strains, the pExpARG plasmid was linearized with StuI and integrated in *RPS1* locus. pExpARG-pEFR3-EFR3 and pExpARG-pYPP1-YPP1 plasmids were constructed by amplification of gDNA using primers with a unique XhoI site at 5’ and a unique NotI site at 3’ ends, with 1 kb upstream and downstream of the respective ORFs. These fragments were subsequently cloned in pExpARG yielding pExpARG-pEFR3-EFR3 and pExpARG-pYPP1-YPP1, respectively. Finally, pExpARG-pEFR3-EFR3 and pExpARG-YPP1-YPP1 were integrated in the *RPS1* locus yielding the recovery strains *efr3Δ/efr3Δ RPS1::EFR3p-EFR3* and *ypp1Δ/ypp1Δ RPS1::YPP1p-YPP1* respectively.

To generate mutants, we first generated a *stt4* DAmP (Decrease Abundance by mRNA Perturbance) allele by integration of the *SAT1* gene (*via* amplification of pFASAT1 (54) with primers CaSATDAmPpS1 and CaSATDAmPmS2) 5’ of the *STT4* stop codon. The remaining *STT4* copy was replaced with *URA3* using a knockout cassette generated from amplification of pGemURA3 (53) with primers CaStt4pKO and CaStt4mKO. As the *stt4Δ/stt4DAmP* strain behaved identical to the wild-type, we next generated a *stt4Δ/stt4Δ* strain (PY4377) by replacing the allele *stt4DAmP* with *HIS1* using a knockout cassette amplified from pGemHIS1 (53) with primers CaStt4pKO and CaSATDAmPmKO. The *URA3* gene of the *stt4Δ/stt4Δ* strain (PY4377) was then replaced by *SAT1* which was amplified from pGFP-Nat (55) using primers CaURApSAT and CaURAmSAT, so that *URA3* could be subsequently integrated at the *RPS1* locus for murine HDC assays. To generate this URA^+^ strain, pExpURA3 (56) was linearized and integrated in *RPS1* locus yielding PY5040, which was subsequently rendered ARG^+^ by transformation with linearized pExpARG plasmid, yielding PY5111. In order to reintegrate *STT4* at the *STT4* locus, we generated a pSTT4-STT4-STT4t cassette in which unique AscI and PmeI sites were inserted into the *STT4* terminator (623 bp 3’ of the stop codon) by site directed mutagenesis using primers CaStt4term_mAscIPmeI and CaStt4term_pAscIPmeI. Subsequently *ARG4* was amplified from pFaARG4 plasmid using primers CaARG4AscI-S1 and CaARG4PmeI-S2, yielding plasmid pExpARG-STT4-STT4-STT4t::ARG4. To generate the recovered strain, the pSTT4-STT4-STT4t::ARG4 fragment (digested by XhoI and NotI) from this plasmid was used to replace *stt4::HIS1* in PY5040, resulting in *stt4Δ/stt4Δ::STT4,* PY5119. To generate a prototroph recovered strain *HIS1* was added back using linearized pGemHIS1 resulting in PY5131. To generate the hypomorph *stt4* mutant which encodes Stt4[G1810D], site directed mutagenesis was carried out with pExpARG-pSTT4-STT4-STT4t::ARG4 using primers CaStt4G1810DpEcoRV and CaStt4G1810DmEcoRV, resulting in pExpARG-STT4p-STT4*-STT4t::ARG4, which was subsequently digested with XhoI and NotI and transformed into PY5040 resulting in PY5757.

Plasmids containing the phospholipid reporters for plasma membrane PI(4)P, Golgi PI(4)P, PI(4,5)P_2_ and PS, pExpARG-pADH1-GFP-(PH^OSH2[H340R]^)_2_-GFP, pExpARG-pADH1-PH^FAPP1[E50A, H54A]^-GFP (13), pExpARG-pADH1-GFP-PH^Plc^*^δ^*-PH^Plc^*^δ^*-GFP (14) and pExpARG-pACT1-GFP-yeLactC2 (36), respectively, were linearized and integrated into the *RPS1* locus. For expressing these reporters in the Stt4[G1810D] strain (PY5757), the *ARG4* gene was replaced by *SAT1* using the primers CaArgExchS1 and CaArgExchS2 and pFaSAT1 (54). To generate fluorescent protein fusions with Stt4, Efr3, and Ypp1 either 3x-mScarlet or mTurquoise2 was amplified using primers CaStt4pXFPS1 and CaStt4mXFPS2, CaEr3pXFPS1 and CaEr3mXFPS2, CaYpp1pXFPS1 and CaYpp1mXFPS2 respectively and plasmids pFA-*3x-mSc-ARG4* (57), pFA-*3x-mSc-CdHIS1* or pFA-*mTurq2-ARG4* (C. Puerner, M. Bassilana, and R. A. Arkowitz, in preparation).

### Southern blot analyses; RT-PCR; qRT-PCR; and chitin staining

For Southern analysis, EcoRV digested gDNA was separated on a 1% agarose gel, transferred to a nylon membrane and fixed by UV crosslinking as described (36). The hybridization probes were generated by PCR using primers (*STT4*: CaStt4p5199 and CaStt4m5543; *URA3*: CaUra3pXhoI and CaURA3m81) and Amersham ECL Direct Nucleic Acid Labelling and Detection System kit (GE Healthcare UK Limited, Little Chalfont Buckinghamshire, UK) following manufacturer’s instructions. For RT-PCR and qRT-PCR, primers used (Genep-TM/Genem-TM) are listed in Table S3 and RNA extraction was carried out using Master Pure yeast RNA extraction purification kit (Epicentre) from budding cells and cells incubated with serum. For RT-PCR Actin amplification 30 cycles were used and 32 cycles were used to amplify *EFR3, YPP1, STT4, PIK1alpha, MSS4, SAC1 and mSCARLET.* qRT-PCR analyses were carried out as previously described (36) with indicated primers (Genep-TM/Genem-TM; Table S3). For chitin staining exponentially growing cells were fixed and stained with 25 µg/ml Calcofluor White solution and imaged using Spinning disk confocal microscope (58).

### Quantitation of cell wall components using flow cytometry

Logarithmically growing strains were stained for exposed β(1,3)-glucan using an anti-β(1,3)-glucan mAb (400-2; Biosupplies, Australia) primary antibody and a donkey anti-mouse IgG (H+L) secondary antibody conjugated to Alexa Fluor 568 (A10037; ThermoFisher, France), essentially as described (28). Antibody dilutions, 1:600, primary and 1:500, secondary were used. For total chitin, mannan and glucan, Calcofluor White (Fluorescent Brightener 28 M2R; Sigma), Concanavalin A-Tetramethylrhodamine (11540176; ThermoFisher, France) and Aniline Blue soluble sodium salt (10656822; ThermoFisher, France), were used at concentrations of 25 µg/mL, 50 µg/mL and 50 µg/mL, respectively. Cells were fixed with 4% PFA in PBS for all analyses and washed prior to staining. Incubation of cells with ConA was for 30 min and cells were subsequently washed. Flow cytometry was carried out on a Cell Analyser BD LSRFortessa Sorp using 355 nm and 561 nm laser lines with Hoechst Blue (515/30 nm) and PE-Texas Red (600 nm LP; 610/20 nm) filters, respectively. Data were obtained from 100,000 gated events per strain, from 3 independent experiments.

### Microscopic analyses

For colony morphology analysis plates were incubated for 3-6 days prior imaging (36). mScarlet and mTurquoise fusions were imaged as described (57, 58) with 17 x 0.5 µm z-sections. Quantitation of plasma membrane and internal mean signals was performed on central z-sections using the Matlab program Hyphal-Polarity (13). Ratios of plasma membrane to internal signals were normalized by the mean wild-type ratio. Plasma membrane fractions of PI(4)P, PI(4,5)P_2_ and PS were calculated as the ratio of plasma membrane signal of total signal which was then normalized to the mean wild-type ratio. To represent these values between 1 and 0, 0.5 was subtracted from the normalized ratio (the Matlab program detects first signal going in from ROI above background, which is the cytoplasm when there is no plasma membrane localization, hence a value of 0.5 is the absence of plasma signal) and then multiplied by 2 so that wild-type plasma membrane fraction is 1 and no plasma membrane signal is 0. Huygens professional software version 18.04 (Scientific-Volume Imaging) was used for deconvolution of image z-stacks using the appropriate settings for the microscope and excitation source. The signal to noise was set to 10, and the background detection was set to auto, unless otherwise stated. All scale bars are 5 µm, unless otherwise indicated.

### Virulence assays

HDC was induced in 10 Balb/C mice (Charles Rivers, Italy) per group by injecting the lateral tail vein with an inoculum of 5 x 10^5^ cells (59). Animal body weight was monitored daily and animals were sacrificed by cervical dislocation when they had lost more than 20% of their weight. OPC was induced in mice that had been immunosuppressed with cortisone acetate using 7-8 mice per strain as previously described (60). A Vectra Polaris Slide Scanner was used to scan histopathology of murine tongue thin sections, stained with periodic acid-Schiff stain.

### Ethics statements

All OPC animal experiments were approved by the Institutional Animal Care and Use Committee (IACUC) of the Lundquist Institute at Harbor-UCLA Medical Center. All HDC animal procedures were approved by the Bioethical Committee and Animal Welfare of the Instituto de Salud Carlos III (CBA2014_PA51) and of the Comunidad de Madrid (PROEX 330/14) and followed the current Spanish legislation (Real Decreto 53/2013) along with Directive 2010/63/EU.

### Statistical analysis

Differences in mean signals, ratios and percentage of filaments were analyzed by *t*-test and survival experiments with mice were analyzed by the Kaplan-Meier method (Log-rank test) with Graph Pad Prism 8 (La Jolla, CA).

### Transmission electron microscopy

Budding cell samples were processed for electron microscopy and images acquired as described (61). Cell wall thickness was measured from electron micrographs at 3-4 locations around the mother cell and averaged.

## Supplemental material

**Figure S1.**
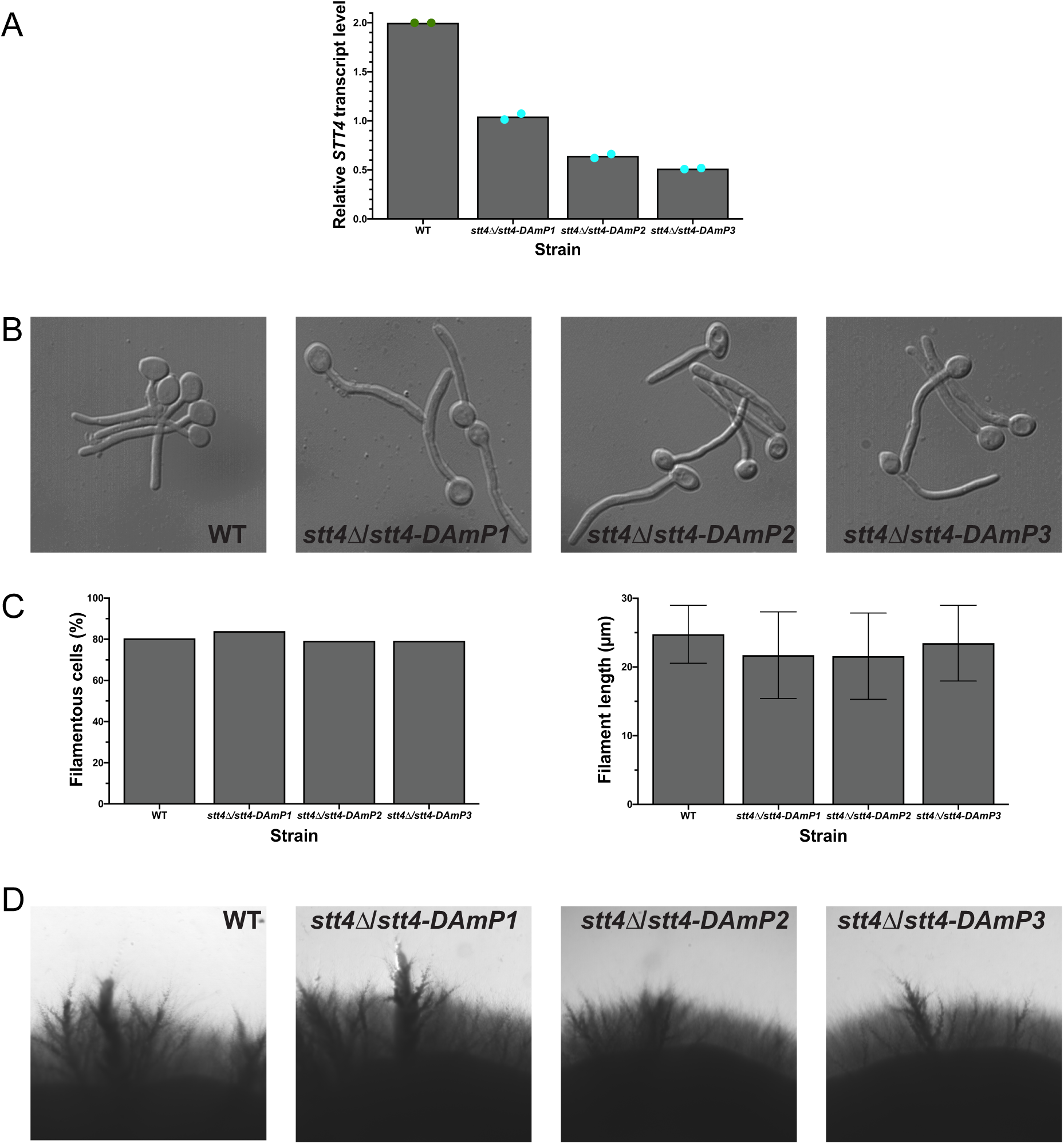
A *stt4* decreased abundance by mRNA perturbation mutants undergoes invasive filamentous growth. A) RT-PCR was carried out on indicated strains (wild-type, PY4861; *stt4*Δ/*stt4-DAmP1-3*, PY4339-4341) using CaSTT4TM1 and CaSTT4TM2 primer pairs. Fragments migrated at the expected sizes. Values are the means of determinations with two primer pairs normalized to *ACT1*. B) Indicated strains were incubated in serum at 37°C for 90 min and images acquired. C) Percentage of filamentous cells (left) (*n* = 350 cells per strain) and hyphal filament lengths (right) were determined (*n* = 60 cells per strain) with error bars indicating SD. D) Indicated strains were incubated for 4 days at 30°C on agar serum plates and images acquired.

**Figure S2.**
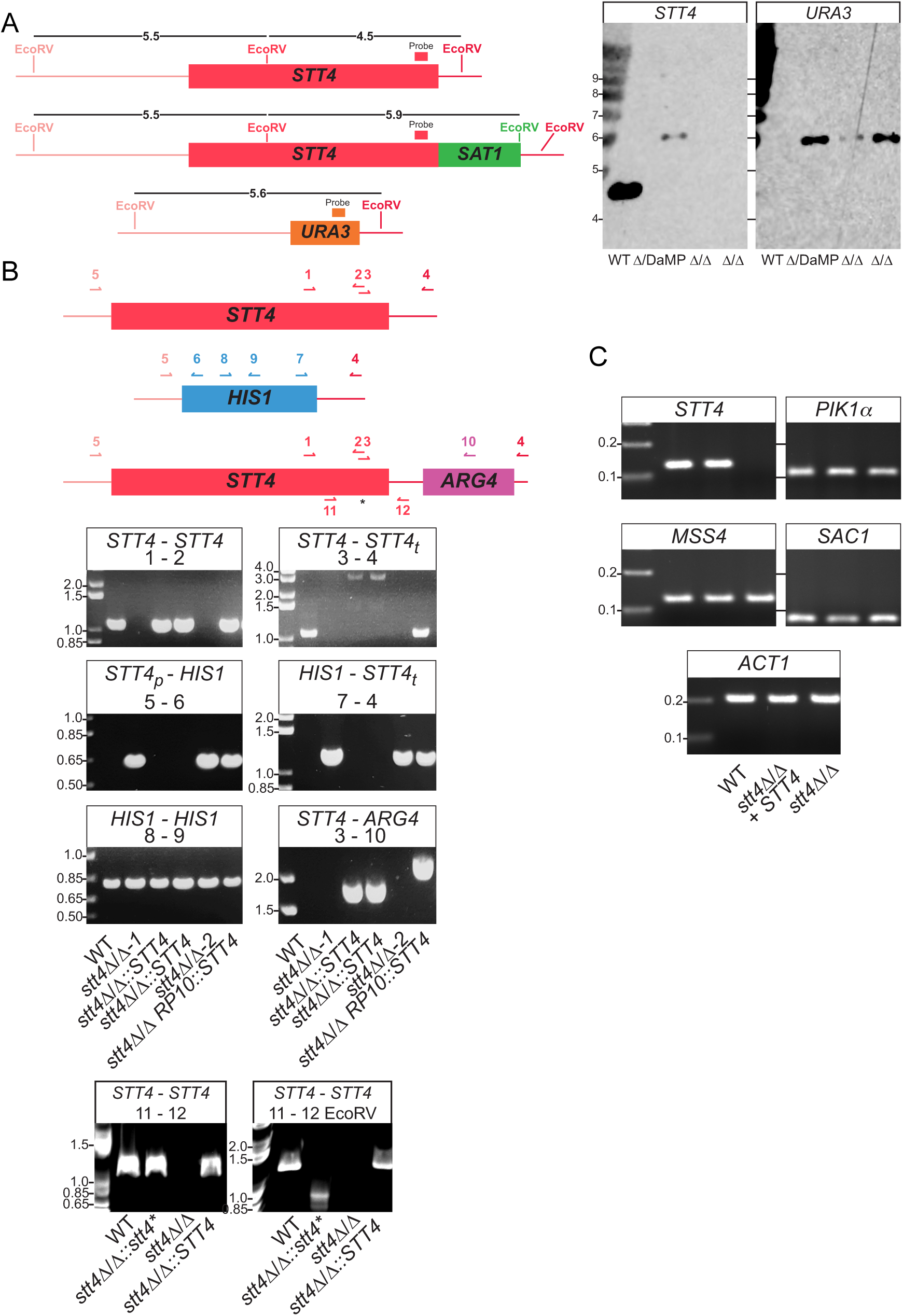
Molecular analyses of *stt4* deletion mutants. A) Southern blot analysis of *stt4* mutants. Schematic representation of chromosomal restriction sites and probes (*STT4* probe, CaStt4p5199 and CaStt4m5543; *URA3* probe, CaUra3pXhoI and CaUra3m81) (left) and Southern blot using *STT4* and *URA3* probes, size in kB, of indicated strains (wild-type, PY4861; *stt4Δ*/*stt4-DaMP*, PY4339; *stt4*Δ/Δ, PY4377 and PY4378) (right) B) PCR analyses of *stt4* mutants. Schematic representation of *STT4* with primers used for strain verification indicated (top) and PCR analyses of indicated strains (wild-type, PY4861; *stt4*Δ/Δ-1, PY5111; *stt4*Δ/Δ::*STT4*, PY5131; *stt4*Δ/Δ-2, PY4414; *stt4*Δ/Δ *RPS1*::*STT4*, PY4433), bottom. Primers, 1 CaStt4p4100; 2 CaStt4m4255; 3 CaStt4p5519; 4 CaStt4m6600NotI; 5 CaStt4pup325; 6 CaHIS1pStop1008; 7 CaHIS1mup152; 8 CaHIS1p214; 9 CaHIS1m836; 10 CaARG4m537; 11 CaStt4p4884; 12 CaStt4m5857. Star indicates position of mutation encoding [G1810D]. C) *PIK1*, *MSS4* and *SAC1* mRNA transcripts are not altered in the *stt4* mutant. RT-PCR was carried out on indicated strains (wild-type, PY4861; *stt4*Δ/Δ + *STT4*, PY5131; *stt4*Δ/Δ, PY5111) with primers for indicated gene amplifications. Fragments migrated at the expected sizes and *ACT1* controls revealed similar amounts of cDNA in each strain.

**Figure S3.**
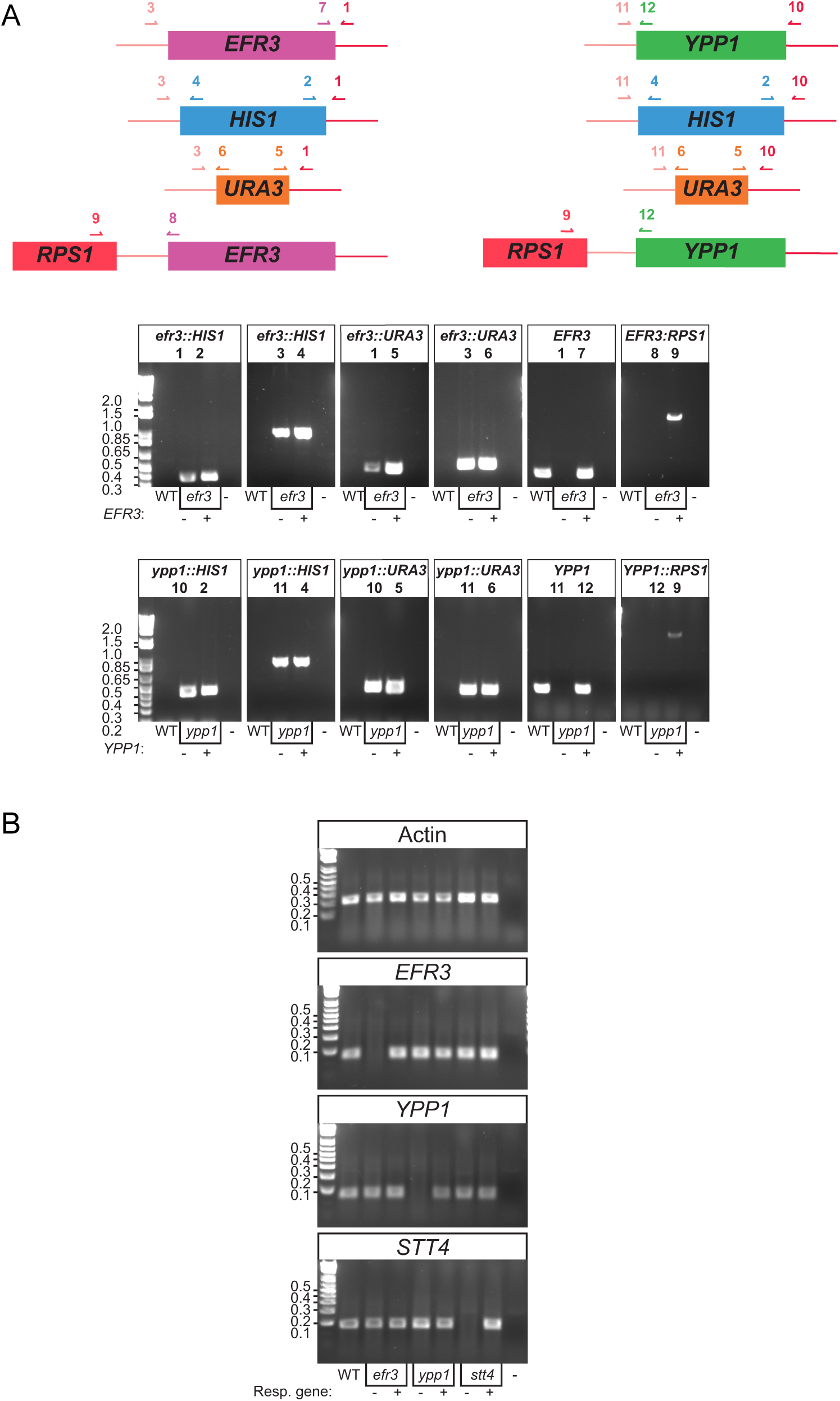
Molecular analyses of *efr3* and *ypp1* deletion mutants. A) PCR analyses of *efr3* and *ypp1* mutants. Schematic representation of *EFR3* and *YPP1* with primers used for strain verification indicated (top) and PCR analyses of indicated strains (wild-type, PY4861; *efr3*Δ/Δ, PY4036; *efr3*Δ/Δ + *EFR3*, PY4039; *ypp1*Δ/Δ, PY4033; *ypp1*Δ/Δ +*YPP1*, PY4040), bottom. Primers, 1 CaYpp1pup167; 2 CaHIS1pStop1008, 3 CaYpp1m3313; 4 CaHIS1mup152; 5 CaURA3p751; 6 CaURA3mup270; 7 CaYpp1m87; 8 CaRPS1p; 9 CaEfr3pup100; 10 CaEfr3m3118; 11 CaEfr3p2692; 12 CaEfr3m127. B) *Efr3*, *ypp1* and *stt4* mutants are only lacking the respective mRNA transcripts. RT-PCR was carried out on indicated strains (same as above and *stt4*Δ/Δ, PY5111; *stt4*Δ/Δ + *STT4*, PY5131) with primers for indicated gene amplifications. Fragments migrated at the expected sizes and *ACT1* controls revealed similar amounts of cDNA in each strain.

**Figure S4.**
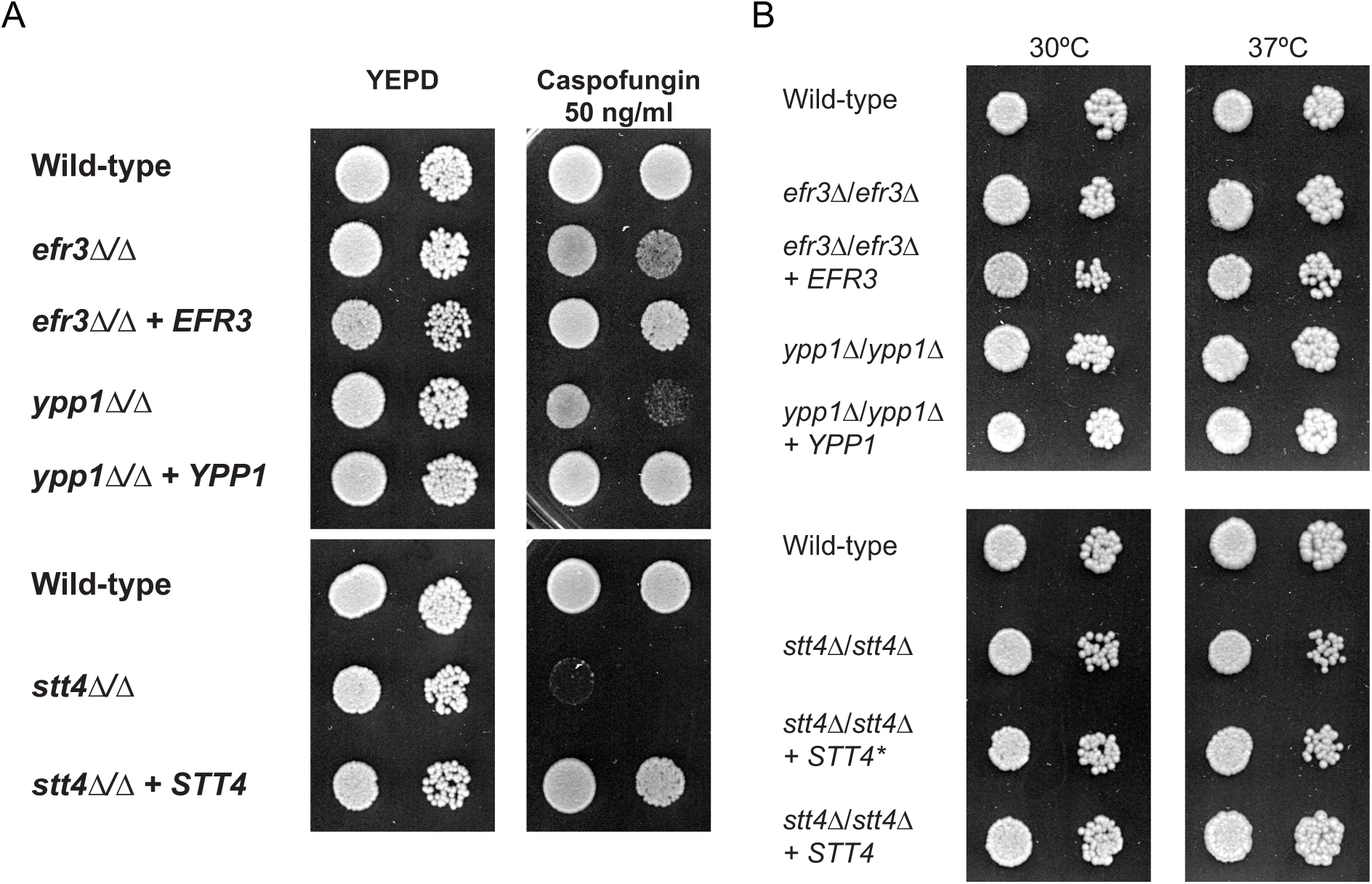
*efr3*, *ypp1* and *stt4* deletion mutants are increasingly sensitive to Caspofungin and grow similar to the wild-type strain at 30°C and 37°C. A) Indicated strains (wild-type, PY4861; *efr3*Δ/Δ, PY4036; *efr3*Δ/Δ + *EFR3*, PY4039; *ypp1*Δ/Δ, PY4033; *ypp1*Δ/Δ + *YPP1*, PY4040; *stt4*Δ/Δ, PY5040; *stt4*Δ/Δ + *STT4*, PY5119) were spotted on YEPD agar containing Caspofungin at the indicated concentration and incubated for 2 days at 30°C. B) Indicated strains (wild-type, PY4861; *efr3*Δ/Δ, PY4036; *efr3*Δ/Δ + *EFR3*, PY4039; *ypp1*Δ/Δ, PY4033; *ypp1*Δ/Δ +*YPP1*, PY4040; *stt4*Δ/Δ, PY5040; *stt4*Δ/Δ + *stt4\** encoding Stt4[G1810D], PY5757; *stt4*Δ/Δ + *STT4*, PY5119) were spotted on YEPD agar and incubated for 2 days at the indicated temperature.

**Figure S5.**
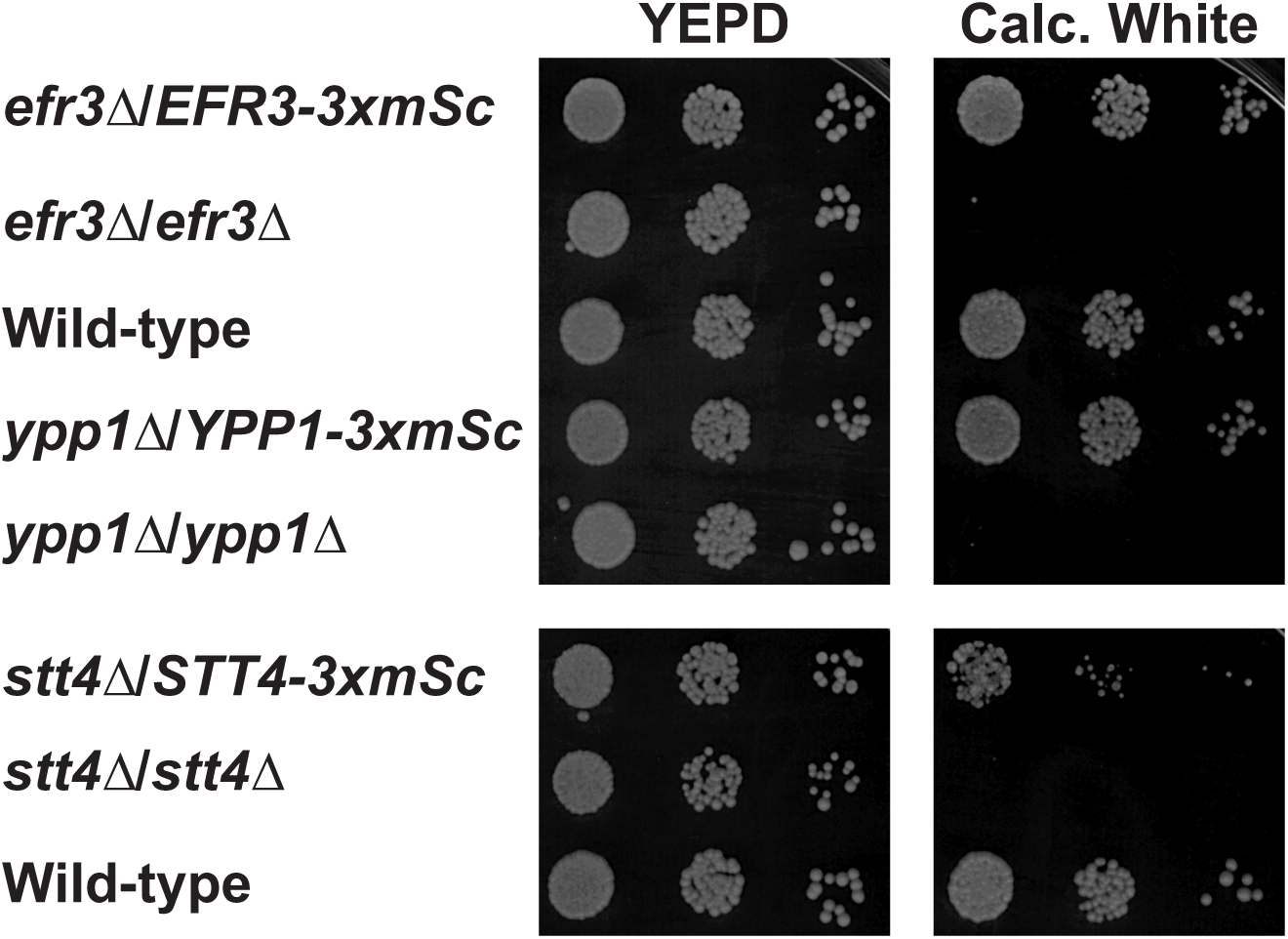
The Efr3, Ypp1 and Stt4 3xmScarlet fusions are functional. The indicated strains (*efr3*Δ/*EFR3-3xmSc*, PY5599; *efr3*Δ/Δ, PY4036; wild-type, PY4861; *ypp1*Δ/*YPP1-3xmSc*, PY5574; *ypp1*Δ/Δ, PY4033; *stt4*Δ/*STT4-3xmSc*, 5601; *stt4*Δ/*stt4*Δ, PY5111) were spotted on YEPD containing Calcofluor White (25 µg/mL) and incubated for 2 days at 30°C.

**Figure S6.**
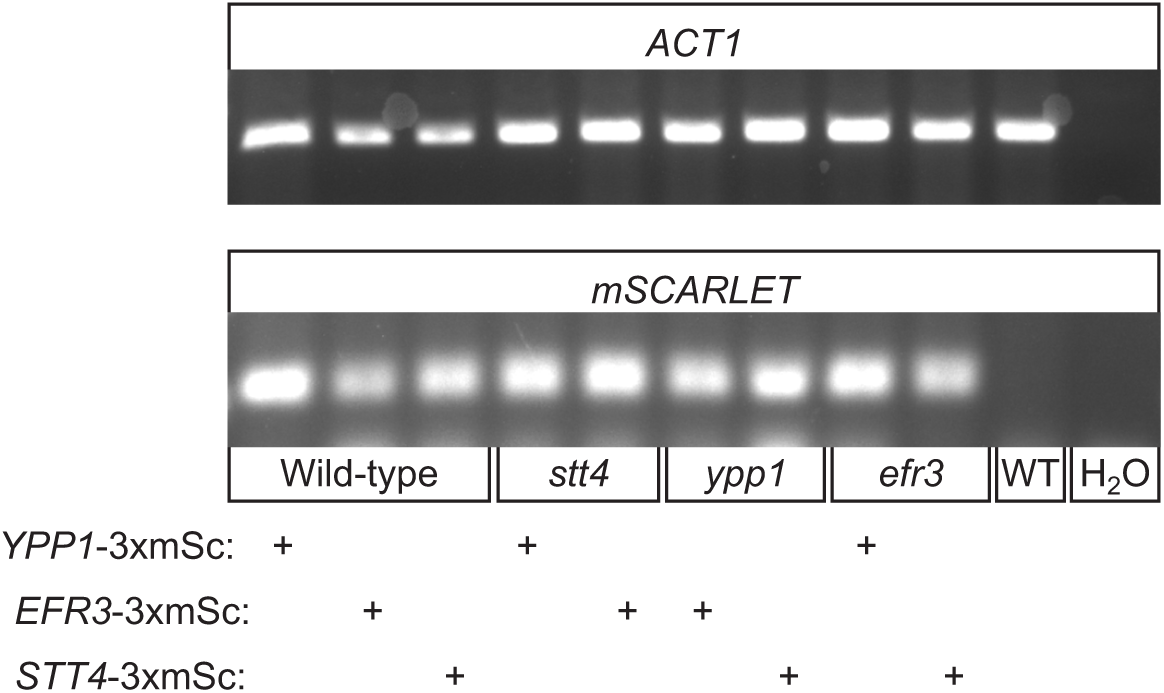
Transcript levels of *EFR1*, *YPP1* and *STT4* mScarlet fusions are not altered in the PI-4-kinase complex mutants. RT-PCR was carried out on the indicated strains (WT, PY6195 Ypp1-3xmSc, PY6197 Efr3-3xmSc, PY6193 Stt4-3xmSc; *stt4* Ypp1-3xmSc, PY6134; *stt4* Efr3-3xmSc, PY6140; *ypp1* efr3-3xmSc, PY6138; *ypp1* Stt4-3xmSc, PY6144; *efr3* Ypp1-3xmSc, PY6136; *efr3* Stt4-3xmSc; WT, PY4860) with primers (CamSCARLETp-TM, CamSCARLETm-TM, CaACT1p-TM and CaACT1m-TM) and for indicated gene amplifications. Fragments migrated at the expected sizes and *ACT1* controls revealed similar amounts of cDNA in each strain. Similar results were observed with two additional mScarlet primer pairs.

**Figure S7.**
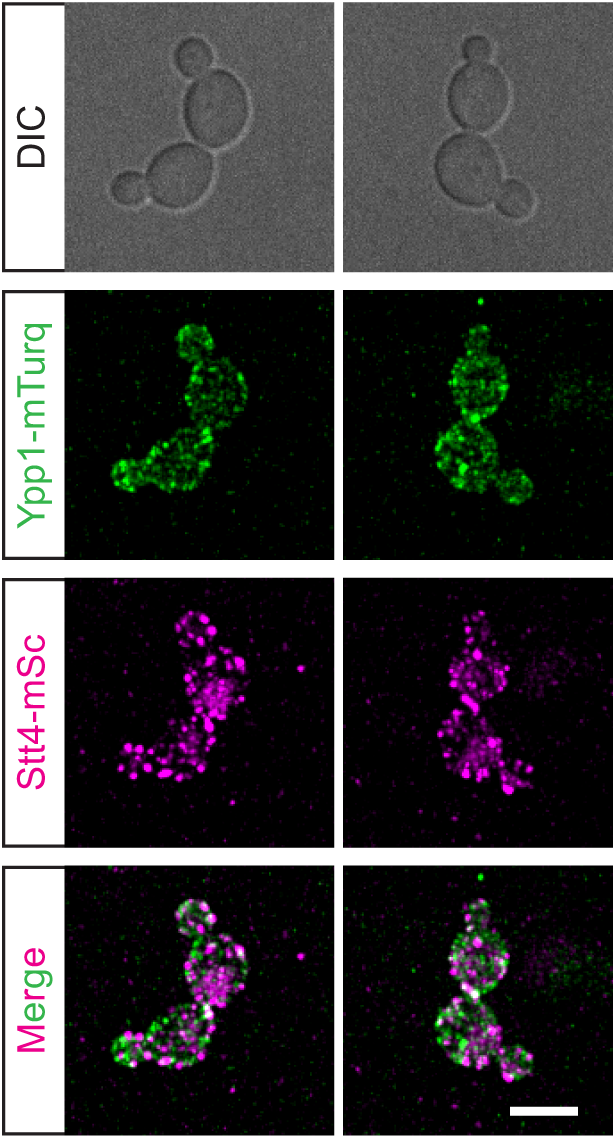
Ypp1 and Stt4 do not colocalize in cortical patches. A strain expressing Stt4-3xmScarlet (magenta) and Ypp1-mTurquoise (green), PY6201, was imaged during budding growth and maximum projections of 17 x 0.5 µm z-sections are shown. Bar is 5 µm.

**Figure S8.**
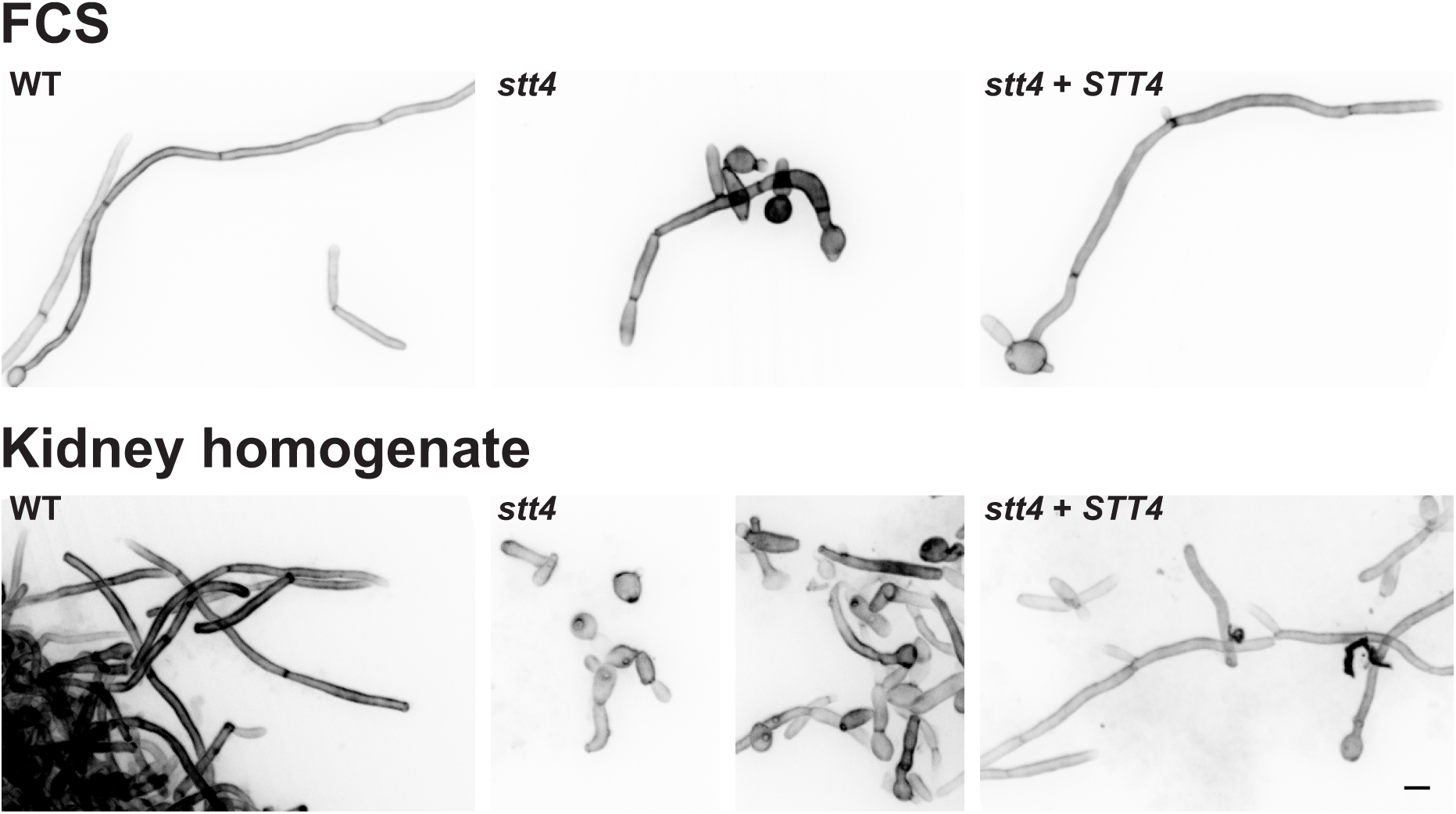
The *stt4* deletion mutant elongates in response to serum and murine kidney homogenates. Indicated strains (wild-type, PY4861; *stt4*Δ/Δ, PY5111; *stt4*Δ/Δ + *STT4*, PY5131) were incubated with either serum or 0.4 g/mL kidney homogenates for 6 Hr at 37°C, samples were fixed, stained with Calcofluor White and images (31 x 0.4 µm z-sections) were acquired. Maximum projections are shown with an inverted LUT. Bar is 5 µm.

**Figure S9.**
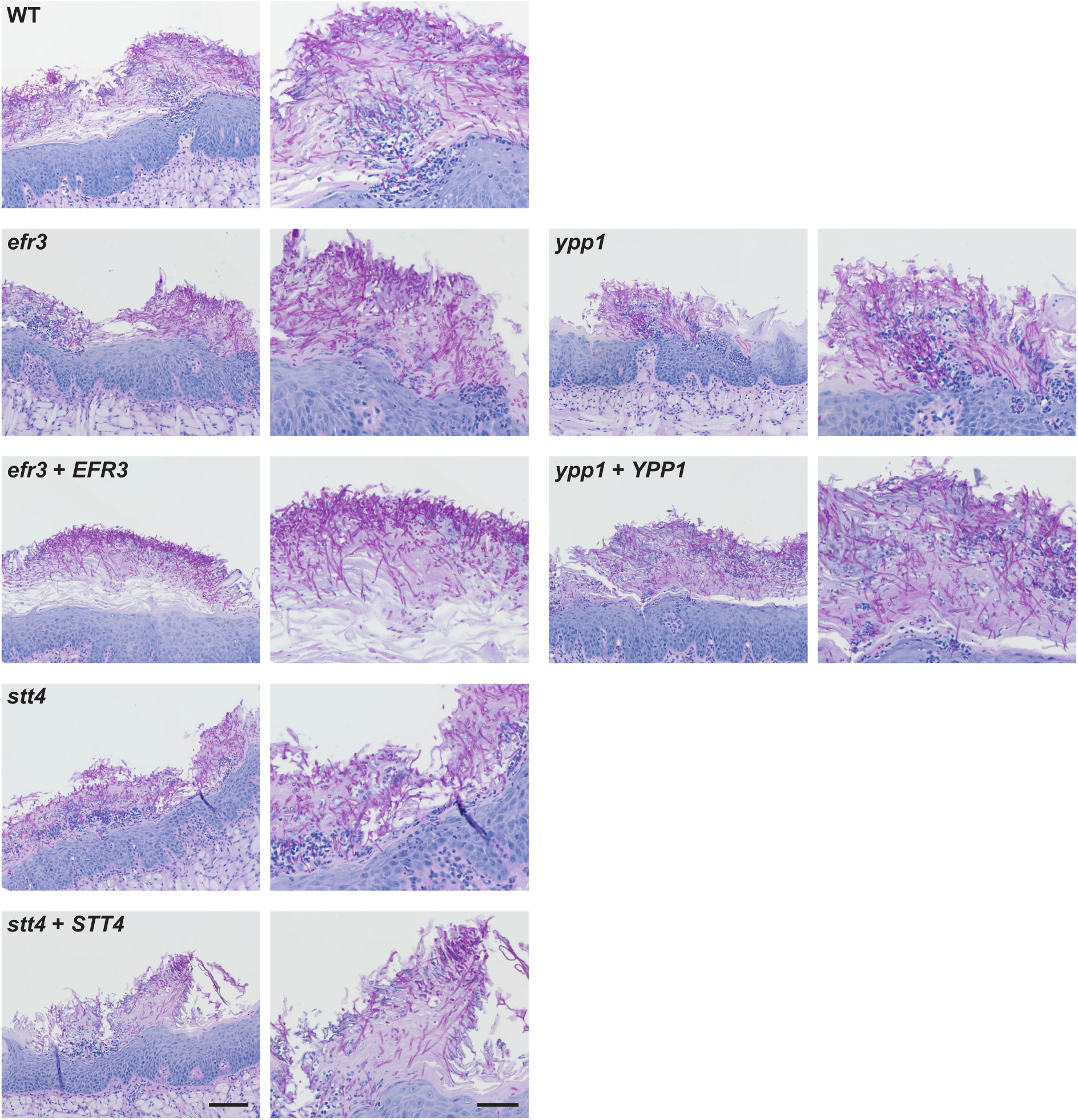
PI-4-kinase complex mutants can filament in a murine OPC model. Histopathology of tongue thin sections from mice infected with indicated strains (see Figure 11C). Thin sections were stained with periodic acid-Schiff stain. Images of regions of infection are shown to highlight fungal morphology, with enlargements of images on left panels (bar, 50 µm) shown on the right panels (bar, 100 µm). Note there were fewer infection sites in mice infected with the *ypp1* mutant, which were also smaller in size.

**Table S1.**
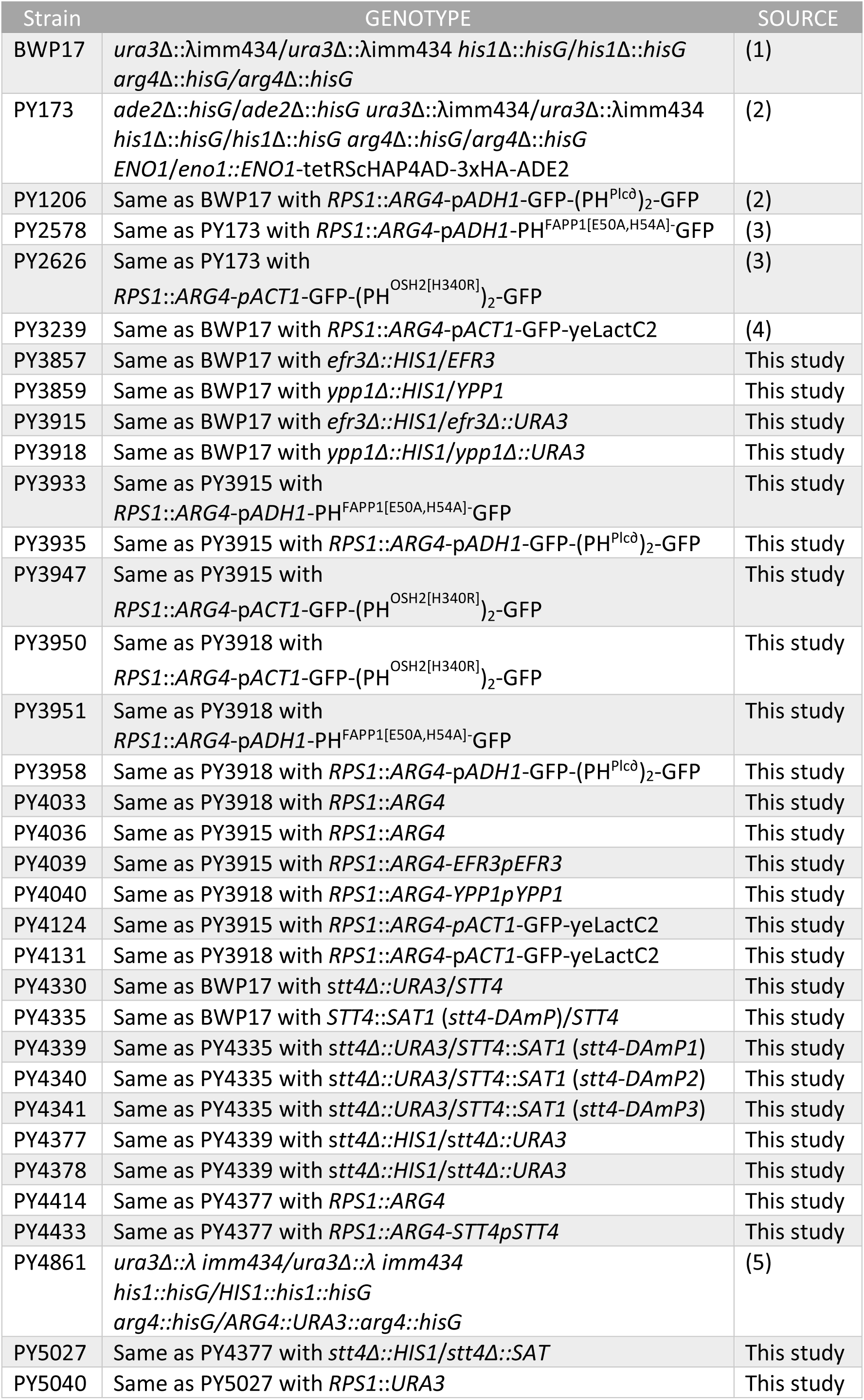
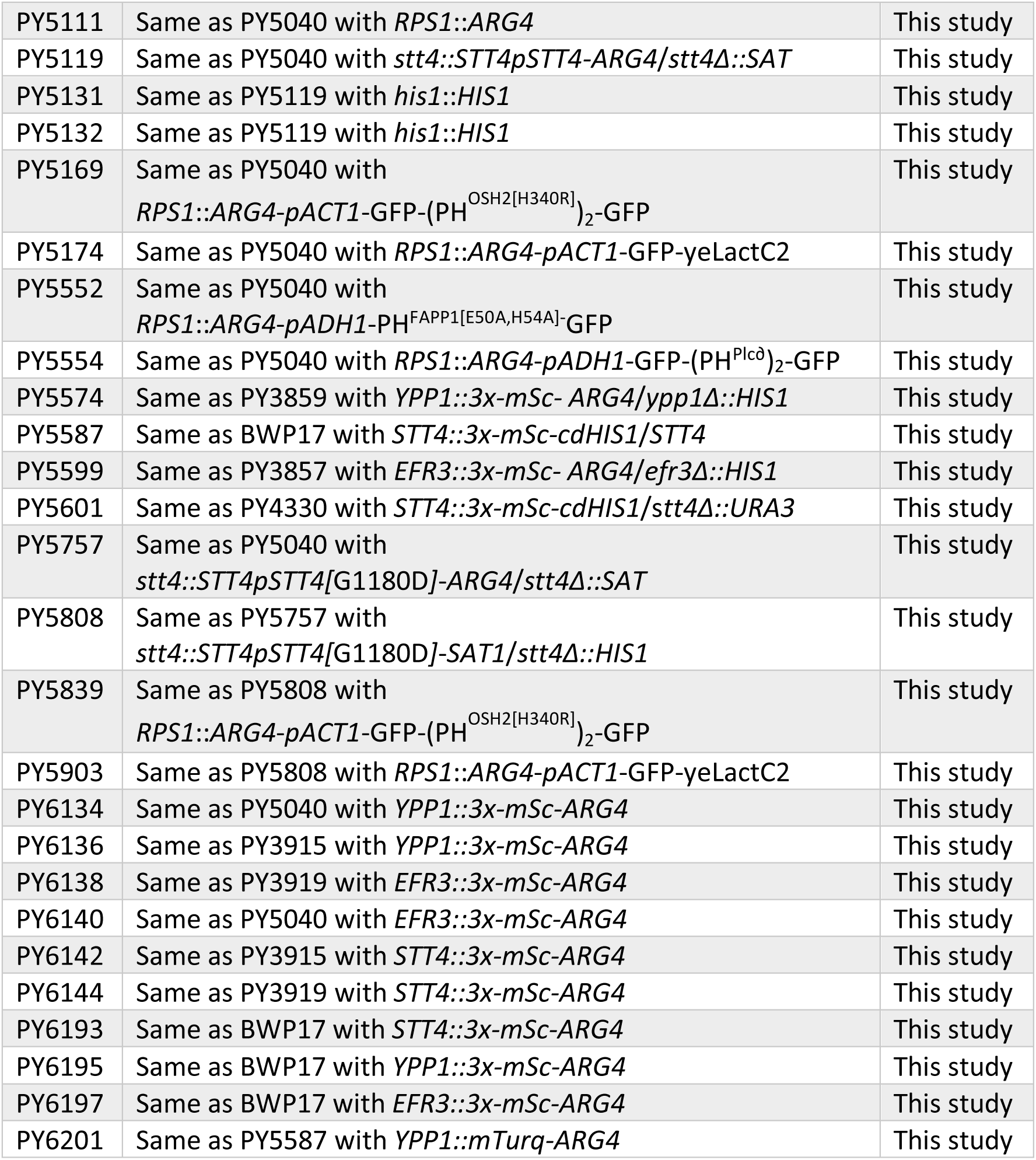
Strains used in this study.

**Table S2.** Plasmids used in this study.

| Plasmid | Source |
| --- | --- |
| pExpARG-pADH1-GFP-(PH<sup>OSH2[H340R]</sup>)<sub>2</sub>-GFP | (3) |
| pExpARG-pADH1-PH<sup>FAPP1[E50A, H54A]</sup>-GFP | (3) |
| pExpARG | (3) |
| pExpARG-pACT1-GFP-yeLactC2 | (4) |
| pExpARG-pADH1-GFP-PH<sup>Plcδ</sup>-PH<sup>Plcδ</sup>-GFP | (2) |
| pGEM-*HIS1* | (1) |
| pGEM-*URA3* | (1) |
| pFA-*SAT1* | (6) |
| pFA-*HIS1* | (6) |
| pFA-*URA3* | (6) |
| pFA-*ARG4* | (6) |
| pFA-3x-*mSc-ARG4* | (7) |
| pFA-3x-*mSc-cdHIS1* | (C. Puerner, M. Bassilana, and R. A. Arkowitz, in preparation) |
| pExpARG4-p*EFR3-EFR3* | This study |
| pExpARG4-p*YPP1-YPP1* | This study |
| pExpARG4-p*STT4-STT4* | (2) |
| pExpARG4-p*STT4-STT4-STT4t::ARG4* | This study |
| pExpARG4-p*STT4-STT4\*-STT4t::ARG4* | This study |
| pFA-*GFPY-NAT* | (8) |
| pFA-*mTurq-ARG4* | (C. Puerner, M. Bassilana, and R. A. Arkowitz, in preparation) |

**Table S3.**
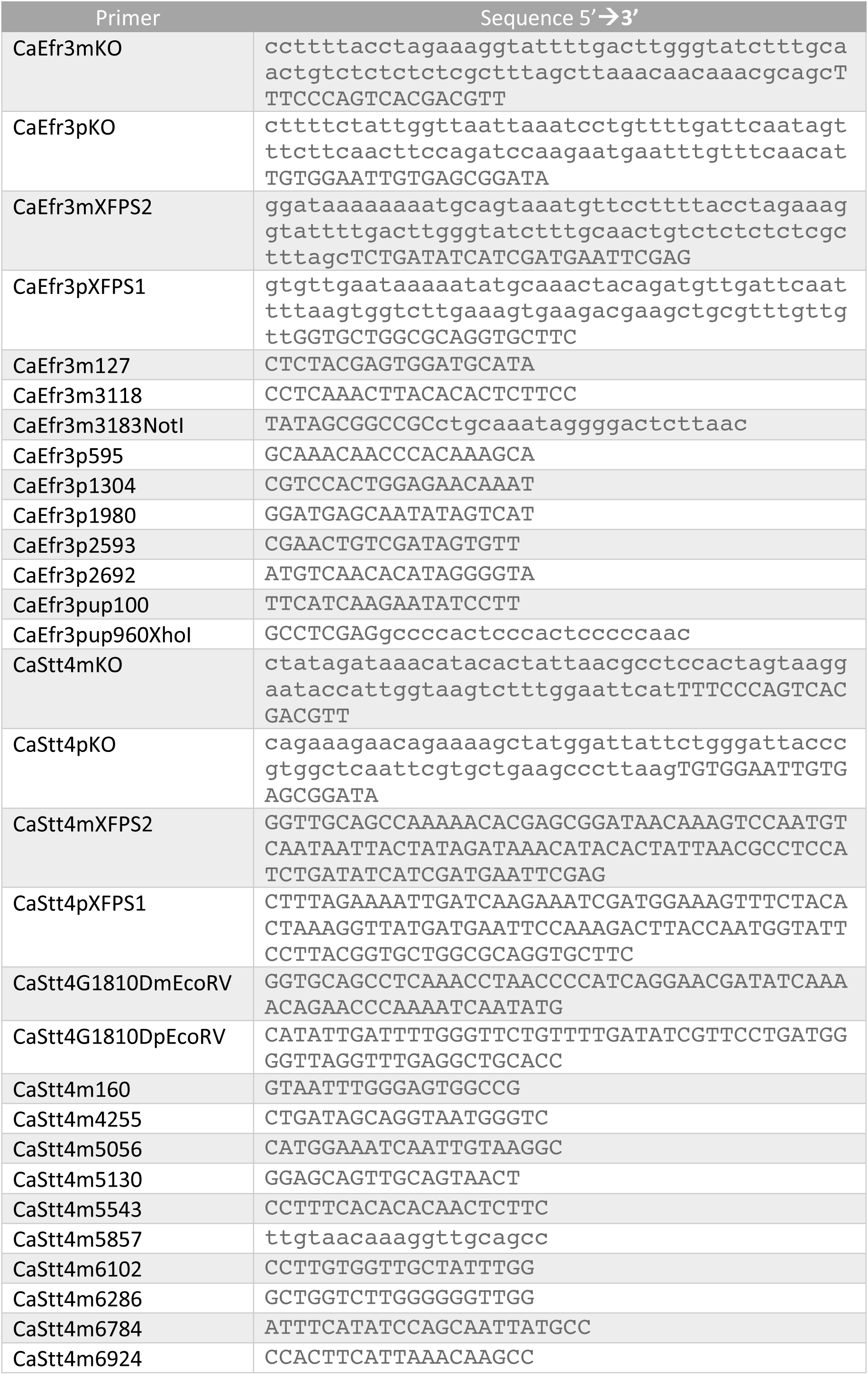
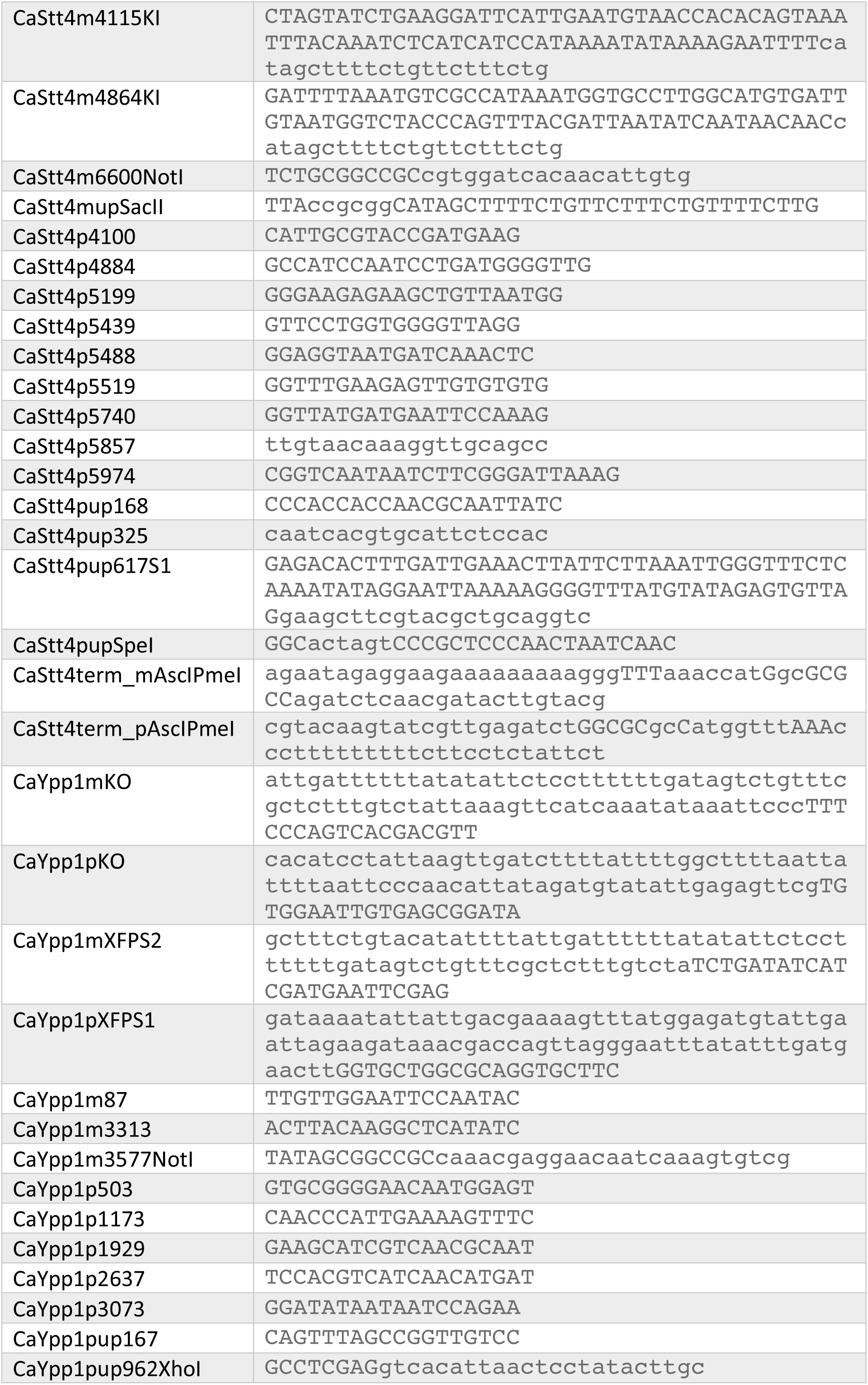
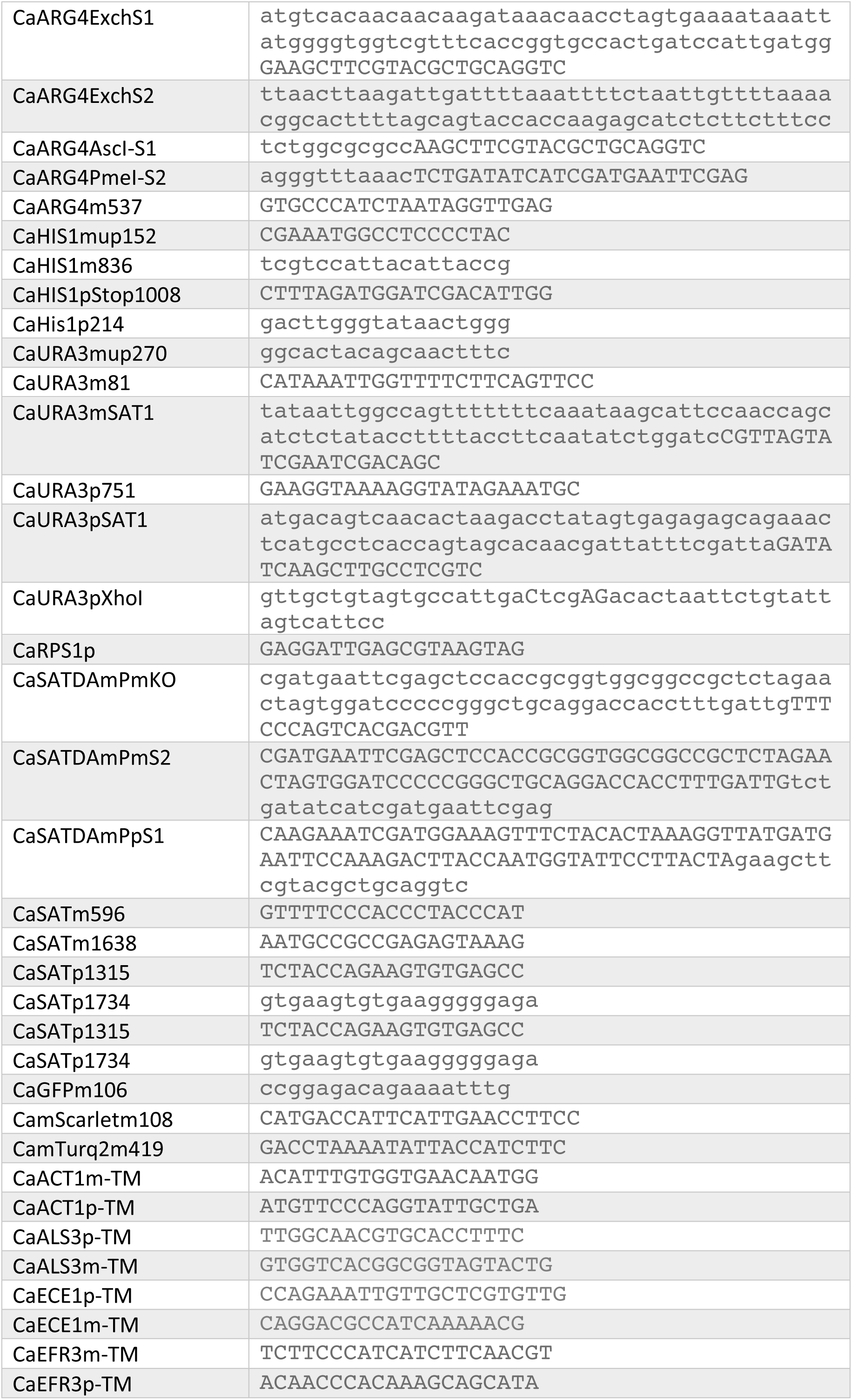
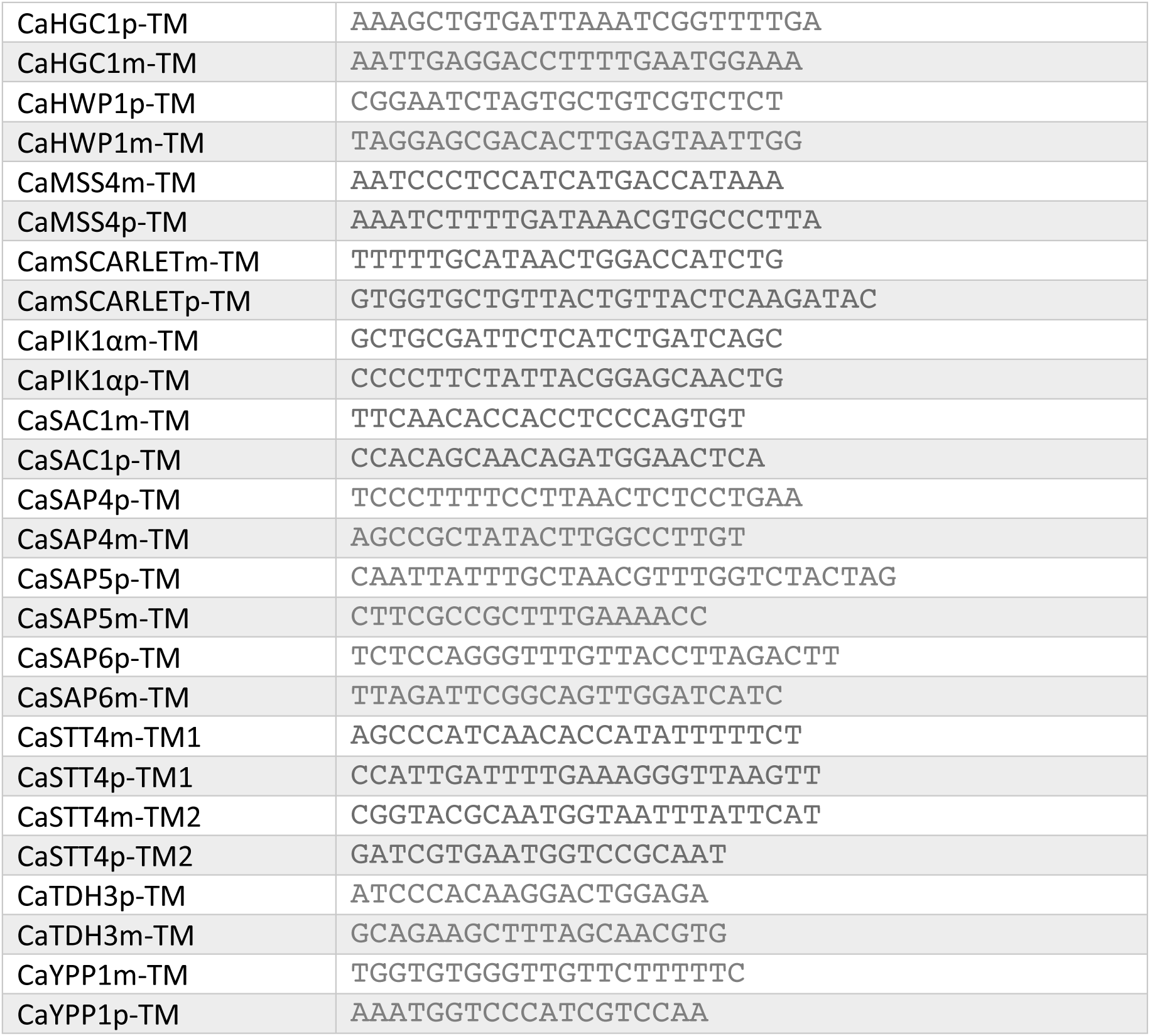
Primers used in this study.

